# Reverse transcribed ssDNA derepresses translation of a retron antiviral protein

**DOI:** 10.1101/2025.10.22.683967

**Authors:** Karen Zhang, Matías Rojas-Montero, Darshini Poola, Josepha M Klas, Arturo Carabias, Dennis J Zhang, Mario R Mestre, Ruiliang Zhao, Ana Dávila-Hidalgo, Guillermo Montoya, Rafael Pinilla-Redondo, Alejandro González-Delgado, Seth L Shipman

## Abstract

Retrons are bacterial immune systems that prevent the spread of phages by initiating a toxic response within infected hosts. All previously characterized retrons produce high levels of multicopy single-stranded DNA (msDNA) in the cell by reverse transcription, which acts as an antitoxin in the absence of phage infection. However, we describe here a non-canonical mechanism for Type VI retrons, which do not produce detectable msDNA in the absence of phage, yet still provide phage defense. Focusing primarily on Retron-Vpa2, a Type VI retron from *Vibrio parahaemolyticus*, we show broad defense against phages and identify triggers of the system within phage recombination systems. Within the Retron-Vpa2 operon, we find a highly enriched, structured transcript that we term a hybrid RNA (hyRNA), which contains both the retron’s reverse transcription template and a translationally repressed toxic effector coding sequence. We find that phage infection induces the accumulation of high levels of msDNA and that this msDNA is necessary for derepressing translation of the antiviral toxin. These findings present key biological and mechanistic insights into a distinct group of retrons while highlighting the diversity of systems that participate in bacterial immunity.

## INTRODUCTION

Bacteria have evolved an arsenal of antiviral defense mechanisms to protect themselves from phages, including a class of defense systems called retrons^1–3^. Retrons operate via abortive infection – upon sensing phage, they release a toxic effector that induces host cell death or dormancy before the phage can propagate, sparing the uninfected bacterial population^2–5^. Retrons are typically three-part systems, consisting of a reverse transcriptase (RT), a noncoding RNA (ncRNA), and a toxic effector^1–3,6^. The RT recognizes a highly structured *msr* region on the ncRNA and reverse transcribes an adjacent *msd* region into multicopy single-stranded DNA (msDNA)^7–13^. The accumulation of high levels of msDNA by ongoing reverse transcription has been considered a hallmark of retrons^12–16^. In fact, the distinctive bands that the msDNA generates on a polyacrylamide gel enabled the discovery of the earliest identified retrons in the 1980s and are used as a signature for Retron-containing species to this day^13–19^.

In canonical retron systems, the msDNA forms a complex with the remaining ncRNA and RT, which together interact with the effector protein as an antitoxin to neutralize its toxic effect^3,20–27^. While toxin-antitoxin motifs are common in phage defense, the incorporation of reverse transcribed DNA in the antitoxin is unique to retrons, enabling these systems to respond rapidly to phage-encoded DNA interacting proteins, such as nucleases^3,28–30^, single-stranded binding proteins^28^, and methyltransferases^3,16,22^. During infection, these phage proteins degrade, perturb, or modify the msDNA, leading to release of the toxin and activation of the defense response^3,16,22,28,29^.

Retrons are a particularly diverse class of defense systems. A bioinformatics study encompassing over 1900 retrons from multiple bacterial phyla classified retrons into thirteen types based on different protein domains found in the highly variable toxic effector component^6^. Following this bioinformatic work, an experimental census was conducted to survey retrons from all thirteen types, including characterization of msDNA production^15^. This census revealed an unexpected trait among a previously untested group: Type VI retrons failed to produce detectable msDNA.

Here, we present the first mechanistic study of Type VI retrons, finding that this type deviates substantially from the canonical mechanism described in previously studied retrons. These retrons defend against phage despite the lack of detectable msDNA at baseline. We reveal that the *msr-msd* and effector coding sequence occupy a contiguous transcript which we call the hybrid RNA (hyRNA). Further characterization of the operon shows that the hyRNA secondary structure and an additional accessory protein result in translational repression of the toxic effector. Counter to all other retrons, reverse transcription of the Type VI msDNA is induced during phage infection, specifically by phage-encoded recombination-associated proteins. Rather than serving as an antitoxin, the Type VI msDNA derepresses the translation of the effector. This inverted triggering mechanism constitutes an intriguing departure from our current understanding of retrons, prompting a reconsideration of the mechanistic range of bacterial immunity.

## RESULTS

### Retron-Vpa2 confers broad phage defense without detectable msDNA

Type VI retrons have a characteristic operon that begins with a predicted ncRNA region containing the *msr-msd*, followed by an ORF encoding a small protein (SP) of unknown function (Fig 1a). Downstream of this is another unknown accessory protein containing a helix-turn-helix (HTH) domain, followed by the retron reverse transcriptase (RT).

**Figure 1.**
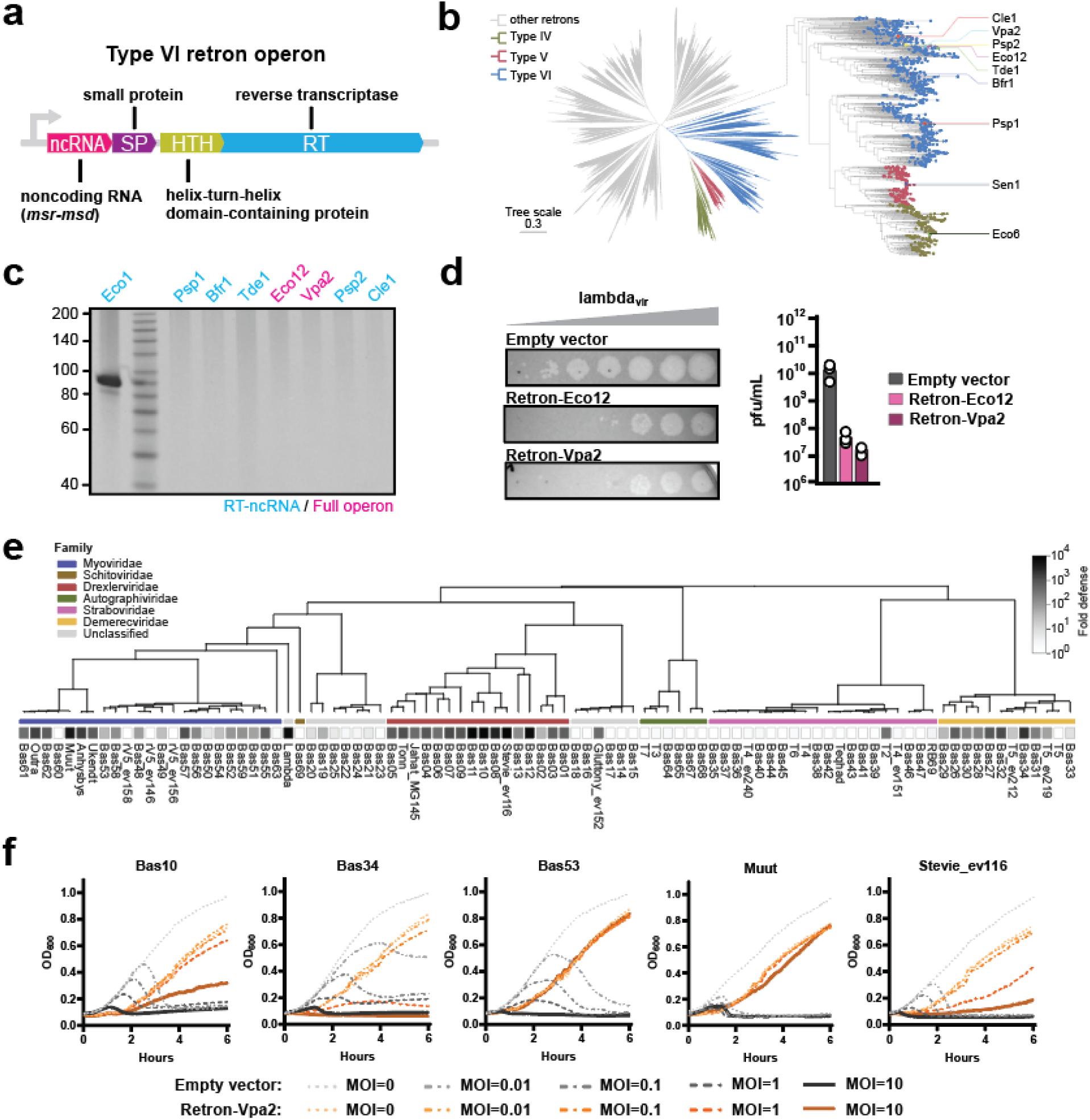
Retron-Vpa2 confers broad phage defense without detectable msDNA. **A)** Schematic of the Type VI retron operon. **B)** Location of Type VI retrons and their close relatives in clade 3 of a retron RT phylogeny. **C)** PAGE analysis for msDNA abundance comparing several Type VI retrons with Retron-Eco1 serving as a positive control for msDNA. A single-stranded DNA ladder with lengths marked in nucleotides is shown for reference. **D)** Spot assay showing titration of phage lambda_vir_ on Retron-Eco12- and -Vpa2-expressing strains of E. coli bSLS.114 relative to an empty vector control. Efficiency of plating is quantified as pfu/mL in the adjacent bar graph (one-way ANOVA P=0.0152, Dunnett’s test corrected for multiple comparisons versus empty vector: Retron-Eco12 P=0.0181; Retron-Vpa2 P=0.0179). **E)** Fold defense conferred by Retron-Vpa2 against a panel of 92 phages, depicted by shaded boxes. Colored bars denote phage family, subfamily, and genus. **F)** Growth curves of E. coli MG1655 expressing either Retron-Vpa2 or an empty vector, infected with five different phages at MOIs of 0, 0.01, 0.1, 1, and 10. Additional statistical details in Supplementary Table 1.

Type VI retrons form a well-supported monophyletic clade (clade 3) within the phylogeny of retron RTs^6^ together with the Type IV (e.g. Retron-Eco6^2^) and Type V retrons (e.g. Retron-Sen1^31,32^). In a previously conducted experimental census of all retron types, we found no msDNA production from Type VI retrons when expressed in *E. coli*^15^. Here, we replicated that result (Fig 1c). Five of the retrons were retested exactly as in the retron census, with expression of just the retron RT and predicted ncRNA (Retrons-Psp1, -Bfr1, -Tde1, -Psp2, and –Cle1). For two of the retrons, we synthesized and expressed the entire retron operon to test whether omission of a component other than the RT and ncRNA in the previous census might have accounted for the lack of msDNA. Of the two retrons tested as a full operon, one had been previously included in the census in the more minimal form (Retron-Vpa2 from *Vibrio parahaemolyticus*), while the other was newly included to check whether matching the host species might yield detectable msDNA (Retron-Eco12 from *E. coli*). Yet, we again observed no msDNA for any Type VI retron tested in any format.

We next tested whether these retrons confer defense against phage infection. We expressed each under their native promoters in an *E. coli* BL21-AI derivative with its endogenous Retron-Eco1 deleted (henceforth bSLS.114)^33^. Despite the lack of msDNA production, we observed robust defense against a virulent strain of phage lambda with each of the Type VI retrons (Fig 1d). For Retron-Vpa2, we then tested for defense in an expanded panel of 92 phages, including the BASEL collection^34^ (Fig 1e). We observed broad defense against the *Demerecviridae*, *Myoviridae*, and *Drexlerviridae* families of phages.

We selected several phages that induced strong defense phenotypes (Bas10, Bas34, Bas53, Muut, and Stevie_ev116) and infected liquid cultures of Retron-Vpa2-expressing *E. coli* MG1655 at a range of MOIs from 0 to 10 (Fig 1f). We observed a clear abortive infection phenotype in Retron-Vpa2-expressing cells for phages Bas10, Bas34, and Stevie_ev116, where the population persists through phage infection in a MOI-dependent manner. For phages Bas53 and Muut, cells expressing the retron grow equally well regardless of MOI. By contrast, cultures without the retron exhibit collapse at every MOI. Overall, these data indicate that Retron-Vpa2 confers protection to the bacterial population in the presence of phage.

### Recombination systems trigger Retron-Vpa2 in two phages

To identify factors responsible for triggering Retron-Vpa2 defense, we examined phage escapees for lambda_vir_ and Stevie_ev116. Whole genome sequencing of the lambda_vir_ escapees revealed various mutations in the single-stranded annealing protein (*beta*) and exonuclease (*exo*) genes of the lambda Red operon, a known recombination system that also includes the RecBCD inhibitor *gam*^35–37^ (Fig 2a). Many of these are early stop and frameshift mutations that result in loss-of-function of the proteins. Of the mutations that do not disrupt the open reading frame, the point mutations in *exo* reside near its catalytic site and may influence its enzymatic activity (Fig S1a-c). The point mutation in *beta* resides in its C-terminal domain which interacts with *exo*^38^ (Fig S1e-f). Stevie_ev116 escapee mutations were primarily found in an exonuclease (*exo*) and recombinase (*rec*) contained in the same operon (Fig 2b, S1d, S1g-h). Due to the presence of an ERF family domain on the recombinase, we predict that this operon is also a recombination system^39^.

**Figure 2.**
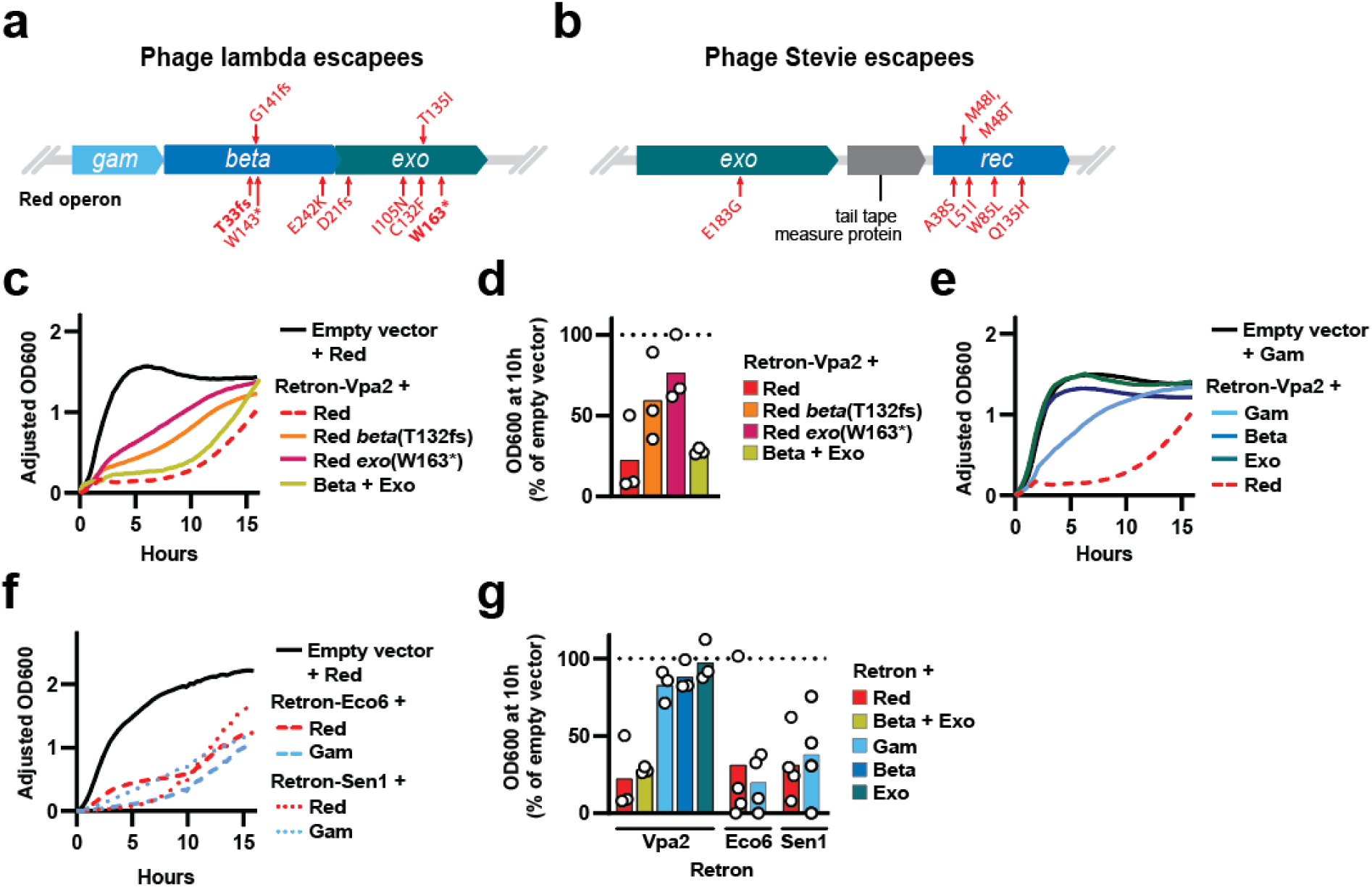
Recombination systems trigger Retron-Vpa2 in two phages. **A)** Operon of the genes mutated in lambda_vir_ escapees of Retron-Vpa2. Specific mutations are labeled in red. Mutations used in trigger assays are in bold. **B)** Operon of the genes mutated in Stevie_ev116 escapees of Retron-Vpa2. Specific mutations are labeled in red. **C)** Trigger assay showing growth curves of liquid cultures expressing Retron-Vpa2 with variants of the lambda Red operon, relative to an empty vector with the Red operon, measured over 16 hrs. Each line is the mean of three biological replicates, error bands are plotted in Fig S2a. **D)** Comparison of OD_600_ measurements from panel (**C**) at the 10 hr timepoint, shown as a percentage of the empty vector control at the same timepoint. Bars show the mean of three biological replicates with individual replicates plotted as white circles (one-way ANOVA P=0.0023, Dunnett’s test corrected for multiple comparisons versus empty vector+Red: Retron-Vpa2+Red P=0.0017, Retron-Vpa2+Beta+Exo P=0.0029, all other conditions ns). **E)** Trigger assay showing growth curves of liquid cultures expressing Retron-Vpa2 with Gam, Beta, or Exo, relative to an empty vector with gam, measured over 16 hrs. Growth curve of Retron-Vpa2 with the Red operon from panel (**C**) shown in a dotted red line for comparison. Each line is the mean of three biological replicates, error bands are plotted in Fig S2b. **F)** Trigger assay showing growth curves of liquid cultures expressing either Retron-Eco6 or Retron-Sen1 with either the Red operon or gam alone, relative to an empty vector with the Red operon, measured over 16 hrs. Each line is the mean of three or more biological replicates, error bands are plotted in Fig S2c. **G)** Comparison of OD_600_ measurements using data from panels (**C**), (**E**) and (**F**) at the 10 hr timepoint, shown as a percentage of the empty vector control at the same timepoint. The Retron-Vpa2 + Red and Retron-Vpa2 + Beta + Exo bars are repeated from (**D**) to permit direct comparison. Bars show the mean of three or more biological replicates with individual replicates plotted as white circles (one-way ANOVA P=0.0002, Šídák’s test corrected for multiple comparisons versus empty vector+Red: Retron-Vpa2+Red P=0.0073, Retron-Vpa2+Beta+Exo P=0.0145, Retron-Eco6+Red P=0.0119, Retron-Eco6+Gam P=0.0027, Retron-Sen1+Red P=0.0118, Retron-Sen1+Gam P=0.0298, all other conditions ns; Šídák’s test corrected for multiple comparisons Retron-Eco6+Red versus Retron-Eco6+Gam and Retron-Sen1+Red versus Retron-Sen1+Gam both ns). Additional statistical details in Supplementary Table 1.

We experimentally validated our candidate trigger genes from lambda_vir_ by co-expressing them with Retron-Vpa2 in liquid cultures of *E. coli* bSLS.114 and measuring OD_600_ over 16 hrs. Co-expression of the retron with the full lambda Red operon resulted in strong growth inhibition compared to a control strain lacking the retron, confirming that this operon is sufficient to trigger the abortive infection phenotype (Fig 2c-d, S2a). We also introduced two mutations that we observed in the escapees (*beta* T132fs and *exo* W163*) to the operon, which each resulted in partial rescue of the growth suppression phenotype. Deleting the third component of the Red operon, *gam*, which is a known trigger of Type IV and V retrons^2,28^, did not rescue growth. Given that the combination of *beta* and *exo* is sufficient to fully trigger the system, we next tested them separately, finding that neither affected cell growth when expressed individually (Fig 2e, S2b). Interestingly, *gam* resulted in slight inhibition when expressed in the absence of *beta* and *exo* (Fig 2e, S2b). This differs from the effect of *gam* on other clade 3 retrons, such as Type IV Retron-Eco6 and Type V Retron-Sen1. For both these retrons, *gam* triggers them to the same degree as the full Red operon, with no requirement for *beta* or *exo* (Fig 2f-g, S2c). This points to a clear distinction between the triggering mechanism of Retron-Vpa2 and that of its close relatives.

### A hybrid RNA encodes a toxic effector and forms a complex with the RT and HTH

We next investigated each component of the Retron-Vpa2 operon. First, we sought to experimentally validate the *msr-msd* region that we predicted based on its secondary structure^15^. Transcripts from the ncRNA region of the retron operon are generally highly enriched in a cell^2^. We therefore performed RNAseq on Retron-Vpa2-expressing *E. coli* bSLS.114 and found that sequencing coverage was enriched not only in the predicted *msr-msd* region, but also in the downstream SP coding sequence (Fig 3a). Performing the experiment with a catalytically dead RT (YADD -> YAAA catalytic site mutation) had no effect on this enrichment (Fig S3). Because this RNA contains both the *msr-msd* and the effector coding region in a contiguous transcript, we decided to name it the hybrid RNA (hyRNA).

**Figure 3.**
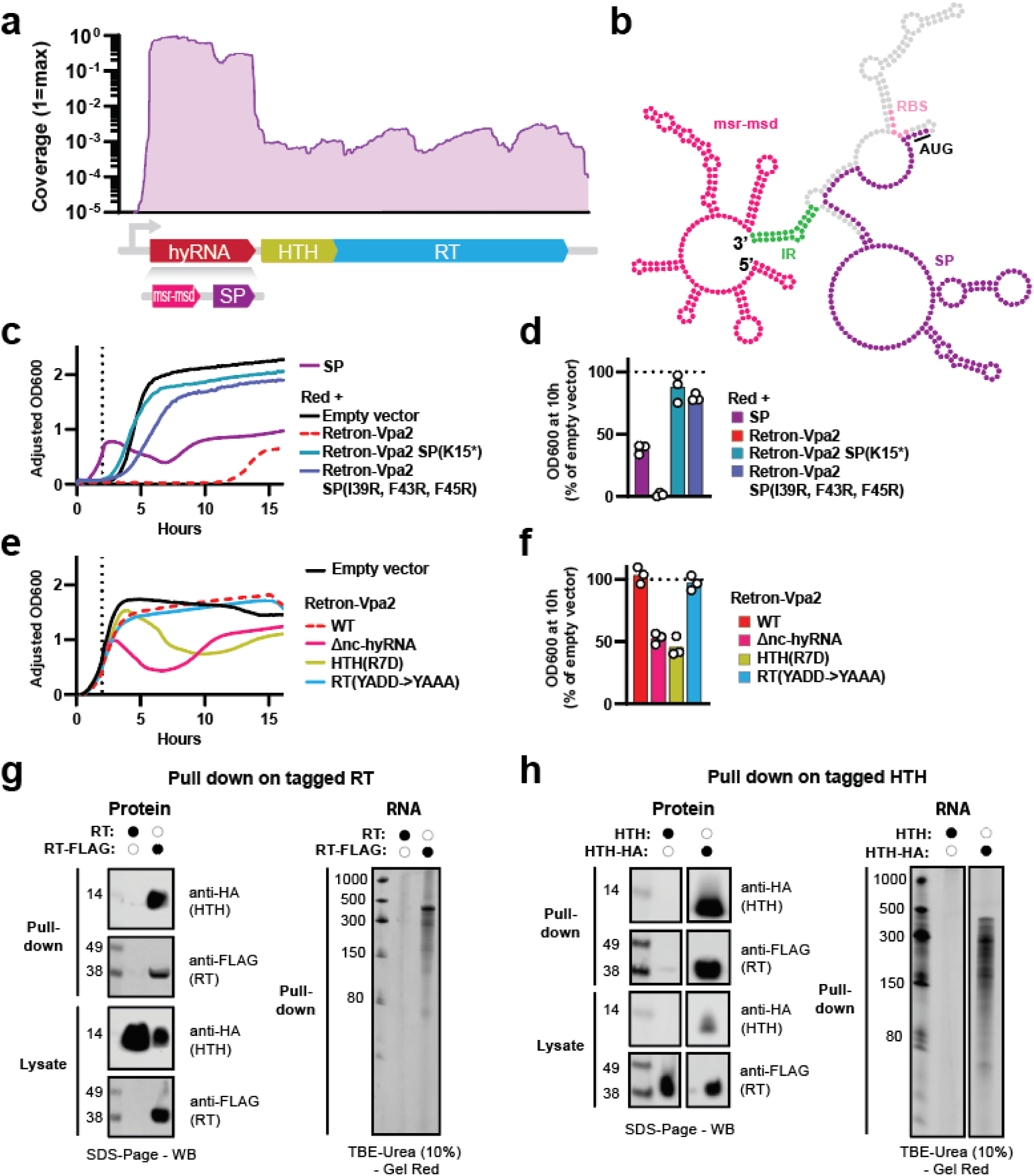
A hybrid RNA encodes a toxic effector and forms a complex with the RT and HTH. **A)** RNAseq coverage (normalized to maximum coverage) of Retron-Vpa2 operon. **B)** Predicted secondary structure of the hyRNA with msr-msd, inverted repeats (IR), ribosome binding site (RBS), SP start codon (AUG) and SP coding region labeled. **C)** Growth curves of liquid cultures expressing SP alone, Retron-Vpa2 triggered with the Red operon, and two Retron-Vpa2 SP mutants triggered with the Red operon, relative to an empty vector with the Red operon, measured over 16 hrs. Cultures were induced at 2 hrs (vertical dotted line). Each line is the mean of three biological replicates, error bands are plotted in Fig S6a. **D)** Comparison of OD_600_ measurements from panel (**C**) at the 10 hr timepoint, shown as a percentage of the empty vector control at the same timepoint. Bars show the mean of three biological replicates with individual replicates plotted as white circles (one-way ANOVA P<0.0001, Dunnett’s test corrected for multiple comparisons versus empty vector+Red: SP P<0.0001, Retron-Vpa2+Red P<0.0001, Retron-Vpa2 SP(I39R, F43R, F45R) + Red P=0.0035, all other conditions ns). **E)** Growth curves of liquid cultures expressing Retron-Vpa2 WT, Δnc-hyRNA mutant, HTH(R7D) mutant, and dRT(YADD->YAAA) mutant, relative to an empty vector, measured over 16 hrs. Cultures were induced at 2 hrs (vertical dotted line). Each line is the mean of three biological replicates, error bands are plotted in Fig S6b. **F)** Comparison of OD_600_ measurements from panel (**E**) at the 10 hr timepoint, shown as a percentage of the empty vector control at the same timepoint. Bars show the mean of three biological replicates with individual replicates plotted as white circles (one-way ANOVA P<0.0001, Dunnett’s test corrected for multiple comparisons versus empty vector: Retron-Vpa2 Δnc-hyRNA P<0.0001, Retron-Vpa2 HTH(R7D) P<0.0001, all other conditions ns). **G)** Anti-FLAG pull-down of cultures expressing Retron-Vpa2, with and without a 3xFLAG-tagged RT. On the left are anti-HA and anti-FLAG Western blots of the pull-down and lysate fractions, with numbers showing protein weight in kDa. On the right is a TBE-Urea PAGE analysis showing detection of hyRNA, with numbers showing length in nt. Complete gels are found in Fig S7a. **H)** Anti-HA pull-down of cultures expressing Vpa2, with and without a HA-tagged HTH. On the left are anti-HA and anti-FLAG Western blots of the pull-down and lysate fractions, with numbers showing protein weight in kDa. On the right is a TBE-Urea PAGE analysis showing detection of hyRNA, with numbers showing length in nt. Complete gels are found in Fig S7b. Additional statistical details in Supplementary Table 1.

The Vpa2 hyRNA is approximately 470 nt in length. Its predicted secondary structure, based on minimal free energy, resembles three arms extending from a central node (Fig 3b). The first arm contains a highly structured region characteristic of a retron *msr-msd*^7,15^. The second arm contains two stem-loops which incorporate the ribosomal binding site (RBS) and start codon of the SP. The third arm contains the remaining coding sequence of the SP. The 3’ end of the transcript then wraps back around and anneals to the first arm via a set of inverted repeats, which resembles the a1/a2 priming region of a canonical retron^9^. A covariance model of the hyRNA independently supports the predicted secondary structure (Fig S4a) and reveals multiple areas of high sequence conservation, including the stems in the first arm, the inverted repeats that link each arm to the central node, the SP RBS and start/stop codons, and a set of direct repeats in the loop regions of the first and second arm (Fig S4b).

Given the SP’s unique position within the hyRNA, we were curious about its role in this system. Its predicted structure reveals a C-terminal domain with a hydrophobic core that is conserved across multiple Type VI homologs (Fig S5a-d). When expressed independently from a synthetic operon, we observed that the SP impairs cell growth at a similar level to triggering the retron with the lambda Red operon (Fig 3c-d, S6a). Introduction of an early stop mutation (K15*) in the SP eliminated the retron’s toxicity upon triggering. Additionally, introduction of three point mutations (I39R, F43R, F45R) in the SP’s conserved hydrophobic core significantly reduced toxicity under the same conditions. From these results, we conclude that the SP alone is the toxic effector of this retron and that its C-terminal domain is essential for toxicity.

We then proceeded to mutate other components of the system in the context of the full operon (Fig 3e-f, S6b). We found that deleting the noncoding region of the hyRNA from the 5’ end up until the RBS (Δnc-hyRNA) was highly toxic. Similarly, mutating a conserved residue in the HTH protein (R7D) was also toxic. However, mutating the RT catalytic domain (YADD -> YAAA)^40^ was not toxic, indicating that reverse transcription is not necessary to neutralize the SP, contrasting with canonical retron mechanisms.

To understand which components of the system physically interact with each other, we next engineered a version of Retron-Vpa2 with FLAG-tagged RT and HA-tagged HTH in a dSP (K15*) background. We expressed the system in *E. coli* MG1655-DE3 and performed a pull-down on either the RT or HTH, followed by a Western blot and TBE-Urea PAGE to detect co-immunoprecipitation of the tagged proteins and hyRNA respectively. Pulling down on the RT led to co-immunoprecipitation of the HTH and hyRNA (Fig 3g, S7a). Pulling down on the HTH led to co-immunoprecipitation of the RT and hyRNA (Fig 3h, S7b). Thus, the RT, HTH, and hyRNA form a complex *in vivo*. We also checked for msDNA in the co-immunoprecipitation but did not detect any in either pull-down (Fig S7c). The co-immunoprecipitated RNA from the RT pull-down was confirmed to be the hyRNA through sequencing (Fig S7d).

### msDNA production is repressed until Retron-Vpa2 is triggered

Given that we detected no abundant msDNA and the catalytic activity of the RT is not necessary for toxin neutralization, we wondered whether the RT could play a non-catalytic role in this retron. We tested phage defense against lambda_vir_ with a version of Retron-Vpa2 carrying a catalytically dead RT and found that inactivating reverse transcription eliminated phage defense (Fig 4a). Given the necessity of reverse transcription for defense, the absence of msDNA was even more puzzling. Thus, we decided to implement a new strategy for detecting msDNA, inspired by our previous work developing a CRISPR-based molecular recorder^33^. We call this strategy Spacer-seq (Fig 4b). Spacer-seq harnesses the *E. coli* Type I-E CRISPR-Cas adaptation machinery, which mediates capture of foreign DNA elements into CRISPR arrays^41–44^. This will “record” the presence of retron msDNA into the host cell’s genome over time. In this CRISPR-Cas system, captured DNA is recorded as 33 nt spacers via the integrase proteins Cas1 and Cas2^43,44^.

**Figure 4.**
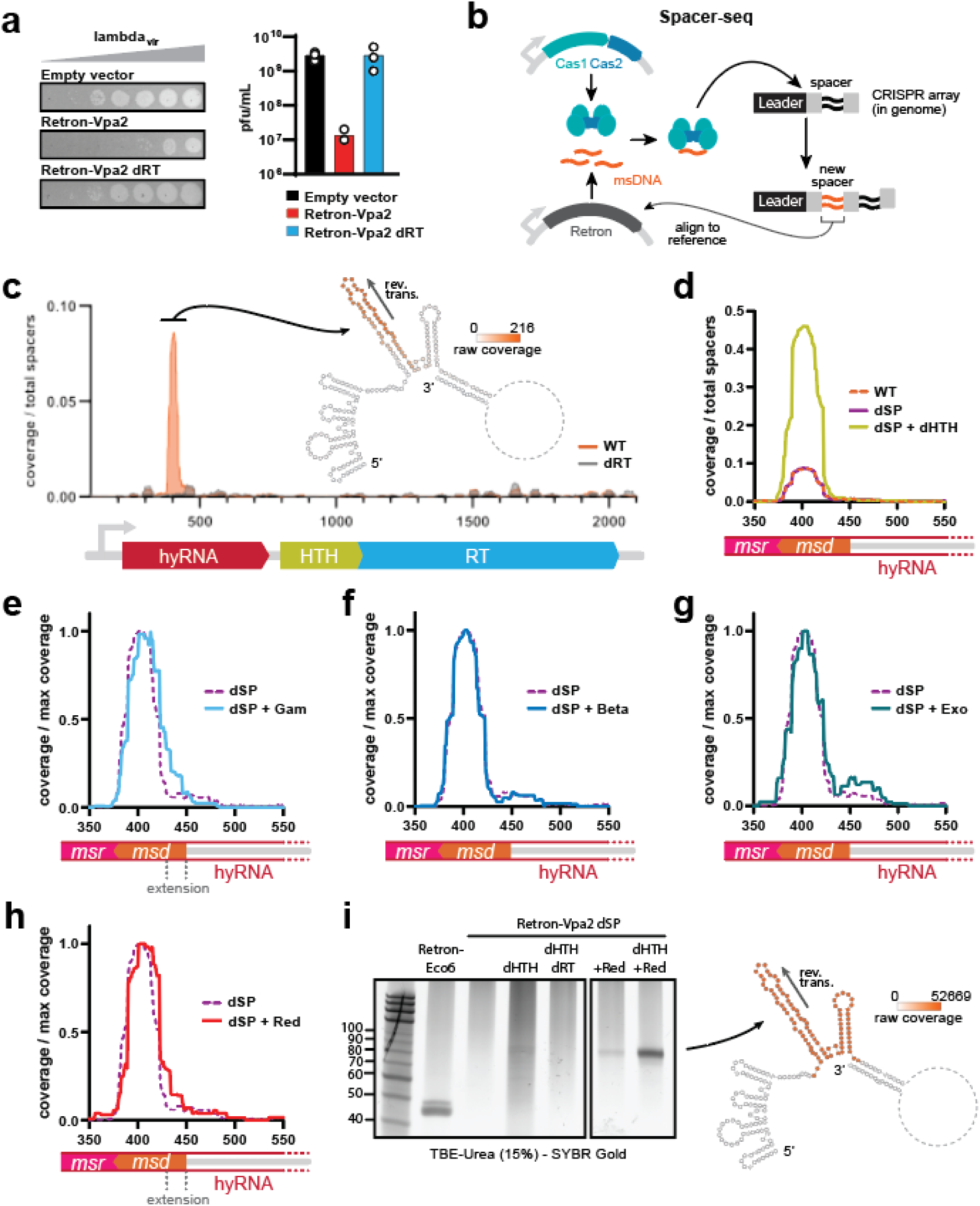
msDNA production is repressed until Retron-Vpa2 is triggered. **A)** Spot assay showing titration of phage lambda_vir_ on Retron-Vpa2 wild-type vs. dRT expressed in E. coli bSLS.114, relative to an empty vector control. Efficiency of plating is quantified as pfu/mL in the adjacent bar graph. **B)** Schematic of Spacer-seq. **C)** Per base coverage (normalized to total spacers acquired) of spacers aligning to the Retron-Vpa2 operon for the wild-type retron compared to a dRT control. Position of aligned spacers is also depicted as raw per base coverage on the predicted secondary structure of the msr-msd region of the hyRNA. Data is a pool of three biological replicates. **D)** Normalized per base coverage of spacers aligning to the msr-msd region of Retron-Vpa2 for various mutants, compared to the wild-type retron from panel (**C**). Data is a pool of three biological replicates. **E-H)** Per base coverage (normalized to maximum coverage) of spacers aligning to the msr-msd region of Retron-Vpa2 for a Retron-Vpa2 dSP mutant with Gam, Beta, Exo, or the Red operon, compared to Retron-Vpa2 dSP alone from panel (**D**). Data is a pool of three biological replicates. **I)** PAGE analysis for msDNA production of Retron-Vpa2 dSP with various mutants, as well as with co-expression of the Red operon. Retron-Eco6 band shown as reference. Ladder on the leftmost lane is single-stranded DNA with increments marked in nucleotides. Complete gel is found in Fig S7b. Position of aligned reads from sequencing the Retron-Vpa2 dSP + Red sample is depicted as raw per base coverage on the predicted structure of the hyRNA msr-msd.

We co-expressed Retron-Vpa2 with Cas1-Cas2 in *E. coli* bSLS.114 (which contains an endogenous CRISPR array but no endogenous Cas proteins). After 24 hrs of induction in liquid culture, we harvested cells and PCR amplified the CRISPR locus. We sequenced these amplicons, identified newly acquired spacer sequences, and mapped them to the Retron-Vpa2 operon. From this, we observed a 63 nt region of enriched coverage (normalized to total spacers acquired) that is absent in a dRT control (Fig 4c). Encouragingly, this region covers a long stem loop in the predicted *msr-msd* portion of the hyRNA. We further validated this method by performing the same experiment on Retron-Eco6 and Retron-Sen1, where we similarly saw enriched coverage of their known *msd* regions (Fig S8a-b). No other peaks of this magnitude were found when mapping coverage across the entire plasmid or *E. coli* genome (Fig S8c). Thus, there is some ongoing msDNA production from Retron-Vpa2, which does not accumulate to a level we can visualize on a polyacrylamide gel.

To identify other factors that might influence msDNA production, we proceeded to perform Spacer-Seq on dSP (K15*) and dHTH mutants of Retron-Vpa2 (Fig 4d). While mutating the SP did not change the distribution of coverage relative to wild-type, mutating the HTH (in a dSP background to avoid toxicity) enhanced coverage by nearly five-fold. We then looked at msDNA coverage of Vpa2 dSP in the presence of different components of the lambda Red operon (Fig 4e-h). Notably, we observed a rightward extension of the spacer distribution peak when Gam is expressed (Fig 4e, h), meaning that this version of the msDNA contains an additional 23 nt at its 5’ end, totaling to a length of 86 nt. This extension also occurs in a Δ*recB* background, indicating that it is linked to Gam’s biological function of RecBCD inhibition^36,37^ (Fig S7a).

Given the increase in Spacer-seq coverage yielded by the dHTH mutation, we attempted once again to visualize msDNA on a gel under similar conditions. Expressing Retron-Vpa2 with a dHTH mutation revealed a series of bands between 60 nt and 90 nt, which are not present in a dRT control or the background dSP strain (Fig 4i, S7b). Additionally, expressing Retron-Vpa2 dSP in the presence of the lambda Red operon yielded a single band between 80 nt and 90 nt, which increases in intensity with a dHTH mutation. Thus, the HTH appears to repress reverse transcription. Sequencing the product with the single band confirmed that it is the same 86 nt product identified in Spacer-seq when Retron-Vpa2 was expressed in the presence of Gam (Fig 4i). We consider this sequence as the full-length msDNA and refer to the 63 nt sequence observed in the absence of Gam as the truncated msDNA.

### SP is translated in the presence of triggers and msDNA

The final step of Retron-Vpa2 phage defense is release of the toxic SP to arrest growth in the infected host. We next aimed to investigate the mechanism of SP neutralization/activation. Because the RBS and start codon of the SP are sequestered within stem loops of the hyRNA, we hypothesized that the toxin is translationally repressed until the retron is triggered. To measure its production *in vivo*, we designed a split GFP complementation assay^45^ where SP abundance is detected through a fluorescence-based readout. In this assay, an optimized superfolded GFP (sfGFP) is split into two fragments which must self-assemble to reconstitute a fluorescent protein. The large fragment contains beta sheets 1-10 (sfGFP1-10), which we express from an independent plasmid. The small fragment contains beta sheet 11 (sfGFP11), which we fuse to the C-terminus of a truncated SP (containing only the first 31 residues) within the Retron-Vpa2 operon. This way, the total length of the hyRNA is preserved and the SP is rendered nontoxic (Fig S10). In *E. coli* bSLS.114, we co-expressed sfGFP1-10 with SP::sfGFP11 in a Δnc-hyRNA mutant of Vpa2 (which we previously saw was toxic with a wild-type SP, thus we expect SP::sfGFP11 to be produced). However, we did not observe fluorescence signal in this condition, nor in a condition where sfGFP11 is not fused to the SP (Fig 5a). We decided to redesign our constructs, this time adding the first 31 residues of the SP to the N-terminus of the sfGFP1-10 as well, with the rationale being that if there existed natural interactions between SP monomers, it would improve the chance of self-assembly. Tagging both sfGFP fragments with part of the SP resulted in a fluorescence signal well over 10000-fold higher than a negative control without sfGFP11 (Fig 5b).

**Figure 5.**
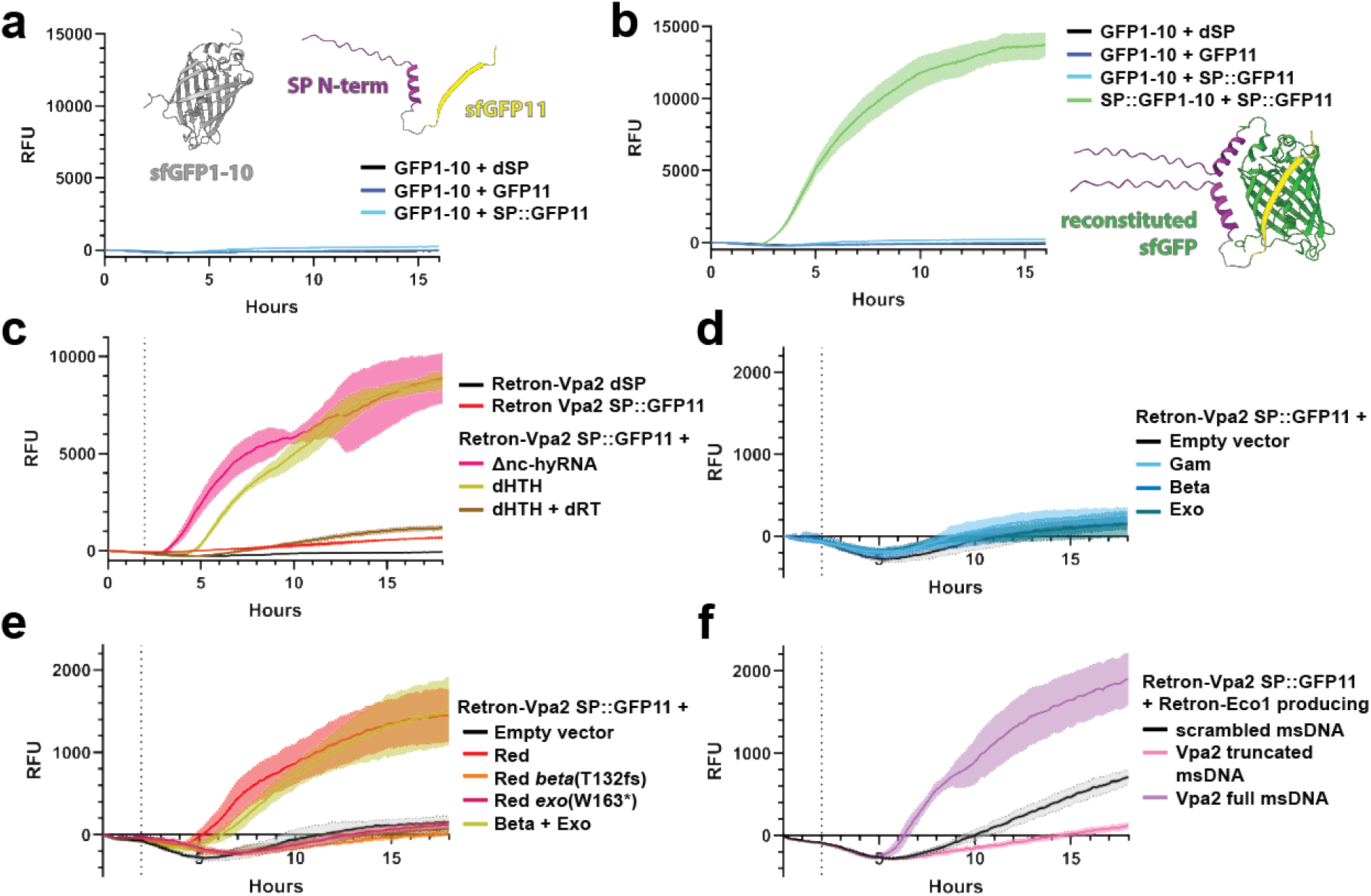
SP is translated in the presence of triggers and msDNA. **A)** Measurement of relative fluorescence units (RFU) for E. coli bSLS.114 liquid cultures expressing a split GFP system over 16 hrs, with induction at 0 hrs. No fluorescence is observed for sfGFP1-10 co-expressed with either sfGFP11 or SP::sfGFP11 (in the context of Retron-Vpa2 Δnc-hyRNA), compared to a control without sfGFP11. Alphafold3-predicted structures for sfGFP1-10 and SP::sfGFP11 are also depicted. **B)** Measurement of RFU for E. coli bSLS.114 liquid cultures expressing a split GFP system over 16 hrs, with induction at 0 hrs. Solid lines indicate the mean and error bands indicate the standard deviation of three biological replicates. Strong fluorescence is observed for SP::sfGFP1-10 co-expressed with SP::sfGFP11 (in the context of Retron-Vpa2 Δnc-hyRNA), compared to the systems tested in panel (**A**). Alphafold3-predicted protein structure shows reconstitution of SP::sfGFP1-10 and SP::sfGFP11. **C)** Measurement of RFU for E. coli bSLS.114 liquid cultures expressing a split GFP system in the context of Retron-Vpa2 Δnc-hyRNA or dHTH mutants over 18 hrs, with induction at 2 hrs (vertical dotted line), compared to a control system lacking sfGFP11. Solid lines indicate the mean and error bands indicate the standard deviation of three biological replicates. **D)** Measurement of RFU for E. coli bSLS.114 liquid cultures expressing a split GFP system in the context of Retron-Vpa2 over 18 hrs, with induction at 2 hrs (vertical dotted line), co-expressed with either Gam, Beta, or Exo, compared to co-expression with an empty vector. Solid lines indicate the mean and error bands indicate the standard deviation of three biological replicates. **E)** Measurement of RFU for E. coli bSLS.114 liquid cultures expressing a split GFP system in the context of Retron-Vpa2 over 18 hrs, with induction at 2 hrs (vertical dotted line), co-expressed with variants of the Red operon, compared to co-expression with an empty vector. Solid lines indicate the mean and error bands indicate the standard deviation of three biological replicates. **F)** Measurement of RFU for E. coli bSLS.114 liquid cultures expressing a split GFP system in the context of Retron-Vpa2 over 18 hrs, with induction at 2 hrs (vertical dotted line), co-expressed with Retron-Eco1 engineered to produce either scrambled msDNA, truncated Retron-Vpa2 msDNA, or full-length Retron-Vpa2 msDNA. Solid lines indicate the mean and error bands indicate the standard deviation of three biological replicates.

We used this assay to measure SP production in the context of a Retron-Vpa2 dHTH mutant and observed high fluorescence (Fig 5c), indicating that the SP is translated when the HTH is inactivated. Since this HTH mutant was previously shown to result in the accumulation of detectable msDNA, we next checked whether SP production in the dHTH background requires msDNA by mutating the RT so that no msDNA can be produced. We found that mutating the RT dramatically reduced fluorescence under these conditions, demonstrating that msDNA is necessary for translation of the SP. Meanwhile, co-expression of the system with variants of the lambda Red operon only resulted in fluorescence in the Red Beta + Exo and wild-type Red conditions, recapitulating our earlier trigger assay results which demonstrated the necessity of *beta* and *exo* for complete retron activation (Fig 5d-e). Interestingly, the maximum fluorescence achieved when triggering the retron with Red was roughly 10-fold less than in the Δnc-hyRNA and dHTH conditions, but still 1000-fold higher than a control where the system is co-expressed with an empty vector. Thus, the SP is translationally repressed, but this repression is alleviated when co-expressing phage triggers, removing the nc-hyRNA, or mutating the HTH, as long as the RT is intact and msDNA can accumulate.

So far, we have observed parallels between the conditions that lead to msDNA production and those that lead to SP production. We wondered whether the two processes are directly linked, where the msDNA molecule itself is derepressing the translation of the SP. To test this, we co-expressed our split GFP system with an engineered version of Retron-Eco1 that produces either a scrambled mDNA sequence, the truncated version of Retron-Vpa2’s msDNA, or the full Retron-Vpa2 msDNA. We observed clear upregulation of SP production in the condition with full Retron-Vpa2 msDNA compared to the scrambled or truncated version (Fig 5f). This shows that abundant Retron-Vpa2 msDNA produced in *trans* from a different retron is sufficient to induce translation of the SP, and that the stem-loop at the 5’ end of the msDNA is essential.

## DISCUSSION

In this work, we uncovered a new mechanism of phage defense by a Type VI retron from *Vibrio parahaemolyticus*, which we named Retron-Vpa2 (Fig 6). This system defends against a broad range of phages, from which we characterized escapees of two phages and identified their recombination-associated trigger genes. This retron, and likely other members of its type, differs from others through its inverted triggering mechanism, where the msDNA acts as the agent for toxin proliferation, rather than inhibition. We determined that Retron-Vpa2 msDNA, which is undetectable at baseline, accumulates 1) in the presence of phage infection, 2) with the expression of phage triggers, or 3) if the HTH protein is mutated. The accumulation of msDNA is necessary for translation of the toxic SP, which is transcribed as part of the hyRNA, but translationally repressed in the absence of msDNA. Finally, the msDNA activates translation of the SP even if produced from another retron in *trans* within the same cell.

**Figure 6.**
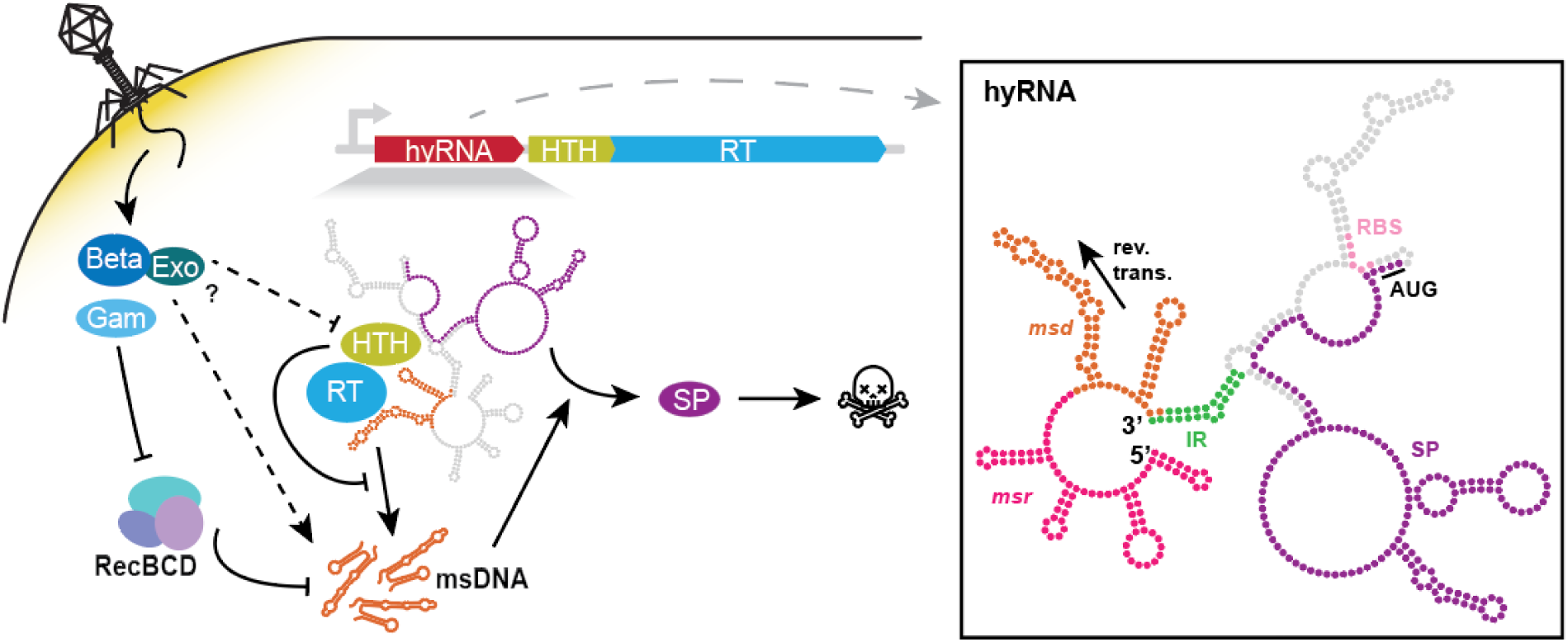
Model of Retron-Vpa2 phage defense mechanism. The core of this system is a complex comprising the RT, HTH, and hyRNA. The hyRNA contains the template for reverse transcription and the coding sequence for SP translation. In its untriggered state, reverse transcription is inhibited by the HTH and the SP is translationally repressed. Phage infection and expression of trigger proteins, such as the Red system of lambda_vir_, induce accumulation of msDNA, which leads to translation of toxic SP.

Given the complexity of this mechanism, there remain several aspects of it that will prompt future investigation. For example, we identify in this study that recombination-associated genes in phage lambda and phage Stevie_ev116 trigger Retron-Vpa2. Still, it is uncertain exactly how this occurs. We hypothesize that the triggers stabilize the msDNA, such that accumulation of msDNA leads to translation of SP. This is supported by the fact that lambda *beta* and Stevie_ev116 *rec* are single-stranded annealing proteins (SSAPs) that bind ssDNA, while *exo* also interacts with DNA^35,39^. Furthermore, it is encouraging that we can identify homologs of these proteins in the majority of phages that Retron-Vpa2 defends against. In the case of lambda phage, the presence of *gam* would also promote msDNA accumulation by inhibiting *E. coli* RecBCD. We think RecBCD, and potentially prophage-associated nucleases, mediate turnover of msDNA from basal levels of reverse transcription. It is also possible that repression of reverse transcription is alleviated by interactions of phage triggers with the HTH.

Through our efforts to understand the role of reverse transcription in this system, we establish Spacer-seq as a discovery-based tool for identifying retron msDNA. Here, it provided evidence that reverse transcription does occur in the absence of triggers, but due to reduced efficiency and/or high turnover rate, msDNA does not accumulate in the cell, and thus cannot be detected through PAGE analysis. We speculate that much of the msDNA captured by the integrases are degradation products of RecBCD and host nucleases, a known source of prespacers for CRISPR adaptation^46,47^. Furthermore, the appearance of the 23 nt 5’ region in the msDNA when RecBCD is inhibited could indicate that the site is normally shielded from the CRISPR integrases by this complex or some other protein, or that it is not synthesized at all.

Perhaps the most intriguing aspect of this retron is its repression of the SP, which requires msDNA production for translation. We think the underlying mechanism involves a type of riboregulator. Notably, the SP RBS and start codon are located within stem loops in the hyRNA, which would sequester them from ribosomal access. In addition, we notice a set of 7 nt direct repeats in the hyRNA that are highly conserved in sequence and structure across the Type VI retrons (Fig S4a-b). One repeat is in a loop region of the *msd* and the other is in a loop region of the predicted riboregulator. When reverse transcribed, the repeat in the *msd* would be complementary to that in the riboregulator, potentially leading to hybridization and subsequent release of the SP RBS and start codon. Importantly, this 7 nt sequence is on the 5’ end of the msDNA, which we demonstrated is specifically necessary for SP translation (Fig 5f). Future work will seek to further understand the specific sequence and structural elements necessary for the msDNA to activate SP translation.

Finally, although the mechanism of SP toxicity remains elusive, we have made some important insights into its biology. We demonstrated that maintaining hydrophobicity of its C-terminal alpha helix is necessary for toxicity (Fig 3b). Furthermore, our results showing augmented fluorescence of split GFP when both fragments are tagged with the first 31 residues of the SP suggest that the N-terminus of the protein mediates oligomerization or colocalization (Fig 5b).

Overall, our study into the biology and mechanism of Retron-Vpa2 reveals a novel retron system where the toxic effector is translationally repressed within a hyRNA transcript and requires phage-induced accumulation of msDNA for expression. These insights will inform future studies on the previously uncharacterized Type VI group of retrons while introducing a new perspective in our understanding of Retron-mediated phage defense.

## METHODS

### Bacterial strains and growth conditions

The *E. coli* strains used in this study were NEB 5-alpha (NEB C2987) for cloning, MG1655 for the large panel phage defense assays, MG1655-DE3 for the pull-down assays (see **Pull-down assays** for details on strain construction), and bSLS.114 (derivative of BL21-AI lacking Retron-Eco1)^48^ for all other experiments. A Δ*recB* derivative of bSLS.114 (bKAZ020) was constructed using lambda Red recombineering^49^ for the experiment shown in Supplementary Figure 9a.

For experiments with liquid cultures, bacteria were grown in lysogeny broth (LB) at 37**°**C, shaking at 250 rpm. Where appropriate, the LB was supplemented with antibiotics and inducers at the following working concentrations: 200 μg/mL ampicillin, 100 μg/mL carbenicillin (GoldBio C-103), 25 μg/mL chloramphenicol (GoldBio C-105), 35 μg/mL kanamycin (GoldBio K-120), 25 μg/mL spectinomycin (GoldBio S-140), 1 mM isopropyl β-d-1-thiogalactopyranoside (IPTG, GoldBio I2481C), and 2 mg/mL L-arabinose (GoldBio A-300).

### Phage strains and propagation

Experiments with phage lambda used a strictly lytic strain (lambda_vir_) that was generously provided by Luciano Marraffini. To propagate the phage, a saturated culture of *E. coli* bSLS.114 was diluted 1:100 in 3 mL LB supplemented with 0.1 mM MnCl_2_ and 5 mM MgCl_2_ (MMB) and grown to an OD_600_ of 0.25. At this point, the culture was infected with phage at a MOI of ∼0.2 and allowed to grow for an additional 16 hrs. After this time, the culture was centrifuged for 10 min at 3,434*g* and the supernatant containing the phage lysate was filtered through a 0.2-μm filter. The titer of the lysate was determined via plaque assay (see **Plaque assays**). Lysates were stored at 4**°**C for further use.

For other phages used in the large panel defense screen, phages were isolated by using sterile inoculation loop to streak from the original stock (obtained from various sources, see Supplementary Table 3) onto a lawn of *E. coli* MG1655 on LB agar. The plate was incubated overnight at 37**°**C. Then, singe plaques were picked and used to infect a 3 mL LB culture of *E. coli* MG1655 at an OD_600_ of ∼0.2. The culture was then grown for ∼5 hrs at 37**°**C. After this time, the culture was clarified via centrifugation and the supernatant containing the phage lysate was filtered through a 0.45 μm filter.

### Plasmid construction

Plasmids containing Retron-Vpa2 (pMRM27) and Retron-Eco12 (pMRM38) were designed to incorporate the wild-type operon (including the native promoter) into a high-copy pET-21 backbone under a T7/lac inducible promoter. These constructs were synthesized by Twist Bioscience. Plasmids containing other retrons tested for msDNA expression were taken from the previous retron census manuscript^15^.

For plasmids containing lambda_vir_ trigger genes, the wild-type trigger genes were PCR amplified from phage lysate and cloned using Gibson assembly (NEB E2621) into a pBR322 backbone under an arabinose-inducible promoter (araBAD). Plasmids containing mutations to the trigger genes were cloned via Q5 site-directed mutagenesis (NEB E0552).

Plasmids containing mutations to Retron-Vpa2 (e.g. Δnc-hyRNA, dRT, dHTH, dSP, dSP) or introducing protein tags to retron components (e.g. HTH-HA, RT-FLAG) were similarly cloned using site-directed mutagenesis.

For Spacer-seq, the plasmid expressing the Cas1-Cas2 genes under a lac promoter (pSCL565) in a pCDF backbone was taken directly from a previous manuscript^50^.

For the split GFP complementation assay, optimized sfGFP1-10 (sequence taken from pAGM22082 in Püllmann et al.^51^, Addgene #153515) was synthesized via Integrated DNA Technologies (IDT) and cloned into a p15a backbone under a lac promoter using Gibson assembly. Site-directed mutagenesis was used to add the first 31 amino acids of the SP to the N-terminus of this plasmid (SP-sfGFP1-10). Site-directed mutagenesis was also used to fuse sfGFP11 (5’-RDHMVLHEYVNAAGIT-3’) with the SP gene (SP::sfGFP11) on a Retron-Vpa2-expressing plasmid. To clone the plasmid used for Retron-Eco1 expression of scrambled msDNA, the retron operon under a T7/lac inducible promoter was amplified from a Retron-Eco1-based editor plasmid (operon is rearranged with the RT preceding the ncRNA and the effector deleted) from a previous manuscript^52^ and cloned into a pCDF backbone using Gibson assembly. Site-directed mutagenesis was used on this plasmid to clone versions that expressed either the truncated or full version of the Retron-Vpa2 msDNA.

All oligos used in this study were synthesized by IDT. Details for all plasmids are listed in Supplementary Table 4.

### Phylogenetic analysis

Homologs of Retron-Vpa2 were identified using the RT amino acid sequence (WP_069548583.1) as a BLASTP search query of the NR protein database (max target sequences = 5000). The resulting sequences were aligned with MAFFT^53^, and a Maximum-Likelihood phylogenetic reconstruction was made with VeryFastTree^54^ with default parameters. Classification of the different retrons was done by annotating the genomic neighborhoods with PADLOC^55^ and used to color the tree.

### msDNA expression and gel analysis

Plasmids containing retrons were transformed into bSLS.114. Individual colonies were picked and grown in 3 mL LB + carbenicillin at 37**°**C until saturation. Each culture was diluted 1:100 in 25 mL LB + carbenicillin, grown for ∼2 hrs at 37**°**C (until OD_600_ reaches ∼0.5), then induced with IPTG and arabinose, after which it was grown for an additional 5 hrs.

DNA was isolated from the cells using the Qiagen Plasmid Plus Midi Kit (Qiagen #12945) and eluted in 150 μL of molecular biology grade water. The eluted DNA was mixed with Novex TBE-Urea Sample Buffer (Invitrogen LC6876) to reach a sample buffer concentration of 1X. This was then heated to 98 C for >=5 min and 15 µL was loaded onto a Novex 15% TBE-Urea gel (Invitrogen EC6885). ss20 DNA Ladder (Simplex Sciences) was also loaded in a separate well as a marker. The gel was run at 200 V for 45 min in preheated (>75**°**C) TBE running buffer, then stained with SYBR Gold Nucleic Acid Gel Stain (Invitrogen S11494) and imaged on a ChemiDoc Imaging System (Bio-Rad).

### Plaque assays

Small-drop plaque assays were performed similarly to Mazzocco et al.^56^, which was used to both titer phage stocks and test for defense. For titering, plaque assays were performed with *E. coli* bSLS.114. For defense experiments with lambda_vir_, a plasmid containing Retron-Vpa2 (or an empty vector control) was first transformed into *E. coli* bSLS.114. Individual colonies were picked and grown in 3 mL LB + carbenicillin until saturation. This culture was then diluted 1:100 in MMB + carbenicillin and grown for 2-3 hrs at 37**°**C. For each plaque assay performed, 200 μL of the passaged culture was mixed with 2 mL of MMB top agar (MMB with 0.75% agar) and poured onto a single well of a rectangular 4-well MMB agar plate (MMB with 1.5% agar). After the top agar has solidified, tenfold serial dilutions in MMB of a phage lysate was spotted onto the plate with 2 μL spots in technical triplicates. After the spots had completely dried, plates placed in an incubator at 37**°**C overnight. Plaque-forming units (pfus) were quantified using the formula: pfu count * dilution factor / mL of lysate spotted = pfu/mL.

For the large panel defense assay, the protocol is similar to above, but plasmids were instead transformed into *E. coli* MG1655 and colonies were grown in LB + ampicillin. Top agar was spiked with 2% of bacterial culture directly from a saturated overnight and phage lysate was spotted onto the plate with 5 μL spots in technical triplicates.

### Phage infection growth curves

A plasmid containing Retron-Vpa2 (or an empty vector control) was transformed into *E. coli* MG1655. Individual colonies were picked and used to start an overnight culture in LB + ampicillin. This saturated culture was then diluted 1:100 into fresh LB + ampicillin and grown to an OD_600_ of ∼0.2. 180 μL of this culture was transferred to a 96-well microtiter plate and infected with phage. All wells received 20 μL of phage lysate that was pre-diluted to various concentrations to achieve final multiplicities of infection (MOIs) of 0, 0.01, 0.1, 1, and 10. The plate was incubated in a plate reader (BioTek Synergy H1 Multimode Reader) at 37**°**C with shaking at 200 rpm for 6 hrs, with OD_600_ measurements taken every 5 min.

### Phage escapee isolation and sequencing

#### Phage lambda_vir_

Isolation of lambda_vir_ escapees generally follows the protocol in Millman et al.^2^. 20 uL of lambda_vir_ lysate (at a titer of ∼10^8^ pfu/mL) was used to infect 200 µL of Retron-Vpa2-expressing *E. coli* bSLS.114 culture at an OD_600_ of 0.3, then incubated for 15 min at room temperature. The infected culture was then mixed into 2 mL of MMB top agar, poured onto a single well of a rectangular 4-well MMB agar plate, and grown overnight at 37**°**C. The next morning, individual plaques were picked and resuspended in 90 μL of phage buffer (50 mM Tris pH 7.4, 100 mM MgCl_2_, 10 mM NaCl). Plaques were left in the buffer for 1 hr at room temperature, with occasional vortexing to release the phage from the agar. This lysate was used to infect a 3 mL MMB + carbenicillin culture of Retron-Vpa2-expressing *E. coli* bSLS.114 at an OD_600_ of ∼0.2, which was grown overnight at 37**°**C. The next morning, the culture was centrifuged at 4347*g* for 10 min and filtered through a 0.2-μm filter. Phage genomic DNA was isolated from 1 mL of this lysate using Norgen’s Phage DNA Isolation Kit (Norgen Biotek 46800).

The purified DNA was prepped for nanopore sequencing on a MinION device using the standard Oxford Nanopore Technologies (ONT) workflow for the Ligation Sequencing Kit (SQK-LSK109) with the Native Barcoding Expansion (EXP-NBD196) on flow cell version R9.4.1 (FLO-MIN106D). The ancestral phage strain was also sequenced alongside the escapees. Sequencing data was base called and aligned to a reference genome (accession J02459.1) using Guppy (v6.1.7 from ONT). Aligned reads were used to generate a consensus sequence for each phage using SAMtools^57^ (v1.18). Escapee-specific mutations were identified through alignment of escapee phage genomes with the ancestral phage genome using Geneious.

#### Phage Stevie_ev116

To isolate Stevie_ev116 escapees, phage lysate was streaked onto a top agar lawn of 2% Retron-Vpa2-expressing *E. coli* MG1655 supplemented with ampicillin. After overnight incubation at 37**°**C, resulting plaques were picked and used to infect a LB + ampicillin culture of Retron-Vpa2-expressing *E. coli* MG1655 at an OD_600_ of ∼0.2. The culture was incubated overnight at 37**°**C to propagate the phage, and phage lysates were harvested the next morning. This process was repeated for three additional rounds to enrich the phage.

Phage DNA was purified from filtered lysates using a combination of enzymatic digestion and silica column-based extraction. To degrade contaminating bacterial nucleic acids, 450 μL of filtered phage lysate was incubated with 50 μL of DNase I 10× buffer, 1 μL of DNase I (1 U/μL), and 1 μL of RNase A. Samples were incubated at 37 °C for 1.5 hours without shaking. Enzymes were subsequently inactivated by adding 20 μL of 0.5 M EDTA. To digest remaining proteins, 1.25 μL of proteinase K was added, and samples were incubated at 56 °C for 1.5 hours without shaking. DNA was extracted using the DNeasy Blood & Tissue Kit (QIAGEN) following a modified protocol. 450 μL of Buffer AL was added to each sample and mixed thoroughly by vortexing, followed by incubation at 56 °C for 10 minutes. Next, 450 μL of 96-100% ethanol was added, and the mixture was vortexed to ensure homogeneity. The entire sample was then transferred to a DNeasy Mini spin column placed in a 2 mL collection tube and centrifuged at approximately 8,000 rpm for 1 minute. The spin column was placed in a new collection tube, 500 μL of Buffer AW1 was added, and centrifugation was repeated at 8,000 rpm for 1 minute. With the column in a new collection tube, another 500 μL of Buffer AW2 was added and samples were centrifuged at 8,000 rpm for 3 minutes. The spin column was then transferred to a clean microcentrifuge tube for DNA elution. DNA was eluted by adding 30 μL of nuclease-free water directly to the center of the membrane, incubating at room temperature for 1 minute, and centrifuging at 8,000 rpm for 1 minute. A second elution was performed by adding an additional 20 μL of nuclease-free water, incubating for 1 minute, and centrifuging as before. Purified DNA was stored at 4 °C until use.

The extracted DNA was quantified using the Qubit dsDNA High Sensitivity Assay Kit (Thermo Fisher Scientific). Sequencing libraries were prepared using the Nextera XT DNA Library Preparation Kit (Illumina), following the manufacturer’s protocol with minor modifications. 1 ng of input DNA per sample was subjected to enzymatic tagmentation, which simultaneously fragmented the DNA and tagged it with adapter sequences. Tagmented DNA was then amplified via limited-cycle PCR using Nextera XT barcoded index primers. Post-PCR clean-up was performed using AMPure XP magnetic beads (Beckman Coulter). Final library concentrations were measured using Qubit and normalized to equimolar concentrations before pooling. Libraries were submitted to Novogene (Beijing, China) for high-throughput sequencing. Sequencing was performed on an Illumina platform using paired-end 2×150 bp chemistry. Adapter trimming and removal of low-quality base calls were performed by Novogene using their in-house quality control pipeline, and demultiplexed reads were returned in FASTQ format for downstream analysis.

### Trigger assays

A plasmid containing the retron being tested (or an empty vector control) was co-transformed with a plasmid containing the trigger gene into *E. coli* bSLS.114. Individual colonies were picked and grown until saturation in 3 mL LB + carbenicillin + chloramphenicol at 37**°**C. Cultures were then diluted 1:100 in 200 uL of fresh LB + carbenicillin + chloramphenicol in a transparent 96-well microtiter plate (Corning 3596). The plate was covered with an air-permeable seal (Breathe-Easy, Diversified Biotech BEM-1) and incubated in a microplate reader (Molecular Devices SpectraMax i3) with shaking at 37**°**C for 16 hrs with OD_600_ measured every 20 min. At the 2 hr timepoint, IPTG and arabinose inducers were added to all cultures. Raw OD_600_ values were adjusted by first multiplying by the empirically determined correction factor of 3.4967, then subtracting the 0 hr measurement for each sample.

### RNA-seq

RNA-seq was performed similar to described in Millman et al.^2^, here using a Retron-Vpa2-expressing strain of *E. coli* bSLS.114. A saturated culture of this strain was diluted 1:100 in 5 mL LB + carbenicillin and grown at 37**°**C to an OD_600_ of 0.6. The culture was centrifuged at 4347*g* for 10 min at 4**°**C. The supernatant was discarded, and the pellet was retained. The pellet was then treated with 100 μL of 2 mg/mL lysozyme solution (in 10 mM Tris, 1 mM EDTA, pH 8.0) and incubated for 15 min at 37**°**C in a 1.5 mL tube. At this time, 1 mL of TRI-reagent (Sigma-Aldrich 93289) was added and mixed by pipetting, then 200 μL of chloroform was added and mixed by pipetting. The sample was incubated at room temperature for 5 min for phase separation. The sample was then centrifuged at 18,213*g* for 30 min at 4**°**C, after which the upper phase was carefully removed and transferred to a fresh tube. This was mixed with 750 μL of cold isopropanol, sodium acetate (NaAOc) to a final concentration of 0.3 M, and linear acrylamide (Invitrogen AM9520) to a final concentration of 10 μg/mL. The sample was frozen either at -20**°**C overnight or -80**°**C for 1 hr, then centrifuged at 18,213*g* for 30 min at 4**°**C. The supernatant was discarded, and the pellet was washed twice with 500 μL of freshly prepared, cold 70% ethanol. For each wash, the ethanol was carefully added without disturbing the pellet, then centrifuged for 15 min at 18,213*g* at 4**°**C, and then the supernatant was discarded. Pellet was air-dried for 5 min, then resuspended in 50 μL of molecular biology grade water and incubated at 56**°**C for 10 min to elute. RNA concentration was quantified via Nanodrop (Thermo Scientific) and Qubit fluorometer (Invitrogen).

15 μg of the eluted RNA was treated with TURBO DNase (Invitrogen AM2238). Then, TRIzol precipitation and washes from the previous paragraph was performed once again on the treated RNA. This time the sample was eluted in 15-20 μL of molecular biology grade water and quantified via TapeStation (Agilent) to ensure a RIN > 5. Ribosomal RNA depletion was performed on 1000 ng of the sample using the Illumina Ribo-Zero rRNA Removal Kit (Illumina 20040526) following the kit protocol and using RNAClean XP beads (Beckman Coulter A63987) for the cleanup steps. Synthesis of cDNA and Illumina TruSeq adapter ligation was performed on 100 ng of depleted RNA using the NEBNext Ultra II Directional RNA Library Prep Kit (NEB E7760) following the kit protocol and using AMPure XP beads (Beckman Coulter A63880) for the cleanup steps. The prepared RNA library was sequenced on an Illumina NextSeq 2000 system. Sequencing reads were aligned to the retron operon using Geneious.

### hyRNA secondary structure prediction

The hyRNA sequence from RNA-seq was input into RNAfold^58^, which predicts secondary structure by computing minimum free energy, and visualized with forna^59^. The resulting structure (see Fig 3b) was independently supported by covariance modeling (see below and Fig S3a).

### hyRNA covariance modeling

Homologs of Retron-Vpa2 were identified using the RT amino acid sequence (WP_069548583.1) as a BLASTP search query of the NR protein database (max target sequences = 100). Flanking nucleotide sequences 1 kb upstream and downstream of RT genes were extracted, clustered at 99.9% sequence identity to remove replicates with CD-HIT^60^ (v4.8.1), and aligned with MAFFT^53^ (v7.525). The resulting alignment was trimmed at the 5′ and 3′ ends to the boundaries of the Retron-Vpa2 hyRNA as determined by RNA-seq. These putative hyRNA homologs were clustered at 95% sequence identity with CD-HIT and realigned with mLocARNA^61^ (v2.0.1). The resulting structure-based multiple sequence alignment was used to build and calibrate a covariance model (CM) of the hyRNA using the Infernal suite^62^ (v1.1.5).

To identify other hyRNA homologs across additional Type VI retron loci using this CM, a database of diverse Type VI retron RTs was generated as previously described (see **Phylogenetic analysis**). Briefly, a broad RT Pfam profile (PF00078) was searched against the ClusteredNR protein database. Hits were retrieved and aligned against the HMM profile using hmmalign –trim from the HMMER suite^63^, and all sequences in the alignment shorter than 150 amino acids were removed with SeqKit^64^. The alignment of remaining RTs (n = 306,351 members) was used to build a tree with VeryFastTree^54^. Tree members were annotated with the best hit from an MMseqs2 easy-search^65^ against a custom reference database of known RTs, revealing a clade of Type VI retron RTs (n = 779 members) that contained Retron-Vpa2.

The 1-kb nucleotide flanking regions of these Type VI retron RTs were extracted and clustered at 95% sequence identity using CD-HIT. The CMsearch function of Infernal was then used to scan the previously built CM through these sequences to identify additional hyRNA homologs, and the final hits (n = 81 Type VI retron loci, including Retron-Vpa2) were evaluated for statistically significant co-varying base pairs with R-scape^66^ (v1.4.0) at a default E-value threshold of 0.05 (Fig S3a). To visualize conserved RNA motifs, a sequence logo was generated from the structure-based multiple sequence alignment of these hyRNA hits using WebLogo^67^ (v3.7.9) (Fig S3b).

### Toxicity assays

A plasmid containing wild-type or mutant Retron-Vpa2 (or an empty vector control) was transformed into *E. coli* bSLS.114. Individual colonies were picked and grown until saturation in 3 mL LB + carbenicillin at 37**°**C. Cultures were then diluted 1:100 in 200 uL of fresh LB + carbenicillin in a transparent 96-well microtiter plate (Corning 3596). The plate was covered with an air-permeable seal (Breathe-Easy, Diversified Biotech BEM-1) and incubated in a microplate reader (Tecan Infinite 200 PRO) with shaking at 37**°**C for 16 hrs with OD_600_ measured every 10 min. At the 2 hr timepoint, IPTG and arabinose inducers were added to all cultures. Raw OD_600_ values were adjusted by first multiplying by the empirically determined correction factor of 6.230707455, then subtracting the 0 hr measurement for each sample.

### Pull-down assays

Three plasmids were constructed containing the Retron-Vpa2 operon with either a C-terminal Flag tag on the RT, a C-terminal HA tag on the HTH, or both tags. For assays using RT as bait, a non-tagged RT plasmid was used as a negative control. Similarly, for assays using HTH as bait, a non-tagged HTH plasmid served as the negative control. A modified *E. coli* MG1655 strain lacking bacterial defense systems and stably integrating the λDE3 prophage (carrying an IPTG-inducible T7 RNA polymerase gene) was generated using the λDE3 Lysogenization Kit (Novagen). This strain is referred to as MG1655-DE3.

Plasmids were transformed into *E. coli* MG1655-DE3 and plated on LB agar supplemented with 100 µg/mL ampicillin. Liquid cultures were grown at 37°C with shaking until reaching an OD₆₀₀ of 0.6. Expression of the operon was induced with 0.2 mM IPTG for 4 hours at 37°C. After induction, OD₆₀₀ was measured and used to normalize sample input across conditions. According to the OD_600_, cells were harvested at >3000 × g for 10 minutes at 4°C and resuspended in 0.8–2 mL of lysis buffer (20 mM Tris-HCl pH 7.5, 150 mM KCl, 1 mM MgCl₂, 1 mM TCEP, 0.1% Triton X-100) to ensure that the total cell concentration is similar in all tubes. Lysis was performed by sonication (3 pulses of 10 seconds at 25% amplitude). Soluble proteins were separated by centrifugation at >16,000 × g for 30 minutes at 4°C. The clarified lysate (800 µL) was incubated with 50 µL of anti-FLAG M2 affinity gel (Sigma-Aldrich, A2220) or anti-HA agarose beads (Thermo, 26181) prewashed with lysis buffer, for 30 minutes at 4°C with gentle agitation. Beads were pelleted at 3000 × g for 3 minutes, and the supernatant was removed. Bound complexes were washed four times with lysis buffer lacking Triton X-100. Beads were resuspended in ∼60 µL of lysis buffer without triton X-100 and divided into three aliquots (∼20 µL each) for protein, RNA, and DNA analysis.

For Protein Analysis by SDS-PAGE and Western Blot, the samples were mixed with NuPAGE™ LDS Sample Buffer (Invitrogen, NP0007) and denatured by boiling for 5–10 minutes at 95°C. Proteins were separated by SDS-PAGE (NuPAGE™ Bis-Tris Mini Protein Gels 4 12–% Acrylamide, Invitrogen, 12030166) and transferred to a nitrocellulose membrane (iBlot™ 3 Transfer Stacks, Invitrogen, IB33001). Entire pull-down samples (40 µL) were loaded, as well as 30 µg of total protein from the input lysate to assess expression levels. The membranes were then blocked for 1 hour at room temperature in TBST (Tris-buffered saline with 0.1% Tween-20) containing 0.5% powdered milk. Detection was performed using HRP-conjugated anti-HA (Anti-HA-Peroxidase, High Affinity, Roche) or anti-FLAG M” antibodies (Sigma, A8592), diluted 1:1000 in TBST with 0.5% powdered milk. Membranes were washed three times for 10 minutes in TBST without milk, and chemiluminescence was visualized using SuperSignal™ West Pico PLUS Chemiluminescent Substrate (Thermo Scientific, 34577).

For RNA analysis, samples were resuspended in urea loading dye (4M Urea, 1 mM Tris pH 7.5, 5 mM EDTA), denatured by boiling for 10 minutes at 95°C, and cooled on ice for 2 minutes. DNA samples were pretreated with ∼ 100 μg/ml of RNAse A for 10 minutes at 37°C before denaturation. RNA and DNA were resolved on 10% TBE-Urea gels (Invitrogen, EC68755BOX) run at ∼180 V for 70–90 minutes. RNA and DNA was visualized by staining with GelRed Nucleic Acid Stain (10000X DMSO, Millipore, SCT122) for 10 minutes prior to imaging.

### Extraction of RNA/DNA from pull-down samples for sequencing

Pull-down samples were prepared as described above. The entire sample (20 µL of beads) was resuspended in 300 µL of nuclease-free ddH₂O and denatured by boiling at 95 °C for 5–10 minutes to release proteins and nucleic acids from the beads. RNA was then purified using the phenol:chloroform extraction method, followed by precipitation with sodium acetate and ethanol. Purified RNA was directly prepped for sequencing using the NEBNext Ultra II Directional RNA Library Prep Kit (NEB E7760) and sequenced on an Illumina NextSeq 2000 system. Sequencing reads were aligned to the reference operon using Geneious.

### Spacer-seq

Spacer-seq experiments were conducted similarly to the naïve CRISPR-Cas adaptation experiments in a previous manuscript^50^. A plasmid containing wild-type or mutant Retron-Vpa2 was co-transformed with the Cas1-Cas2-expressing plasmid into *E. coli* bSLS.114. For experiments involving trigger genes, a third plasmid containing the trigger gene was also transformed. Three colonies (biological replicates) for each sample were picked and grown until saturation in 0.5 mL LB + carbenicillin + spectinomycin (+ chloramphenicol when the trigger-containing plasmid is present) in a 96-well deep well plate (Thermo Scientific 260251) covered with an air-permeable seal (Breathe-Easier, Diversified Biotech BERM-2000) at 37**°**C with shaking at 1000 rpm. Cultures were then diluted 1:100 in 0.5 mL LB + antibiotics + IPTG + arabinose in a new deep well plate and grown for 24 hrs at 37**°**C. Cells were then harvested by boiling 25 μL of culture with 25 μL of molecular biology grade water at 95**°**C for 10 min. 0.5 μL of each boiled lysate was then used as template for PCR amplification of the CRISPR array. Specifically, the primers used in this PCR (Supplementary Table 5) amplified the region between the leader sequence of the array and the second pre-existing spacer. The primers also introduced Illumina TruSeq adapters to the amplicons. Three versions of the forward primer were used, each with a different barcode assigned to a different biological replicate, enabling pooling of replicates for each sample prior to sequencing. Amplicons were then prepared for Illumina sequencing using standard workflows and sequenced on a NextSeq 2000 system.

To analyze sequencing data, Illumina FASTQ files containing sequencing reads for each sample were demultiplexed using the custom barcodes introduced to each biological replicate. Demultiplexed files were then processed using custom code (https://github.com/Shipman-Lab/Spacer-Seq) developed in a previous manuscript^33^, which handles read trimming and extraction of spacer sequences. From this, a FASTA file of all newly acquired spacers was obtained for each biological replicate of each sample. These were again pooled into a single FASTA file for each sample and aligned using Bowtie^68^ (v1.3.1) to a dictionary of references containing the Retron-expressing plasmid, the Cas1-Cas2-expressing plasmid, and the *E. coli* BL21-AI genome. Reads aligning to multiple references were discarded. Coverage per nucleotide position was determined from aligned reads using SAMtools^57^ (v1.18) and normalized either to the total number of newly acquired spacers for a given sample, or to the maximum coverage observed in the Retron-expressing plasmid. Data analysis was performed on Jupyter Lab^69^ (v4.0.7). Visualization of spacer coverage on the Retron-Vpa2 hyRNA secondary structure was performed using VARNA^70^ (v3-93). Visualization of spacer coverage across the full plasmid and genome as shown in Fig S6c was performed using Circos^71^ (v0.69-9).

### Direct msDNA sequencing

Following a protocol from a previous publication^15^, 79 μL of eluted DNA from a bacterial culture midiprep (see **msDNA expression and gel analysis**) was treated with 10 μL of DBR1 (50 ng/μL in-house prep, see previous manuscript^23^) and RNaseH (NEB M0297) in 10 μL of rCutSmart buffer (NEB B6004) and molecular biology grade water to reach a total reaction volume of 100 μL. The reaction was incubated at 37**°**C for 30 min. Then, the product was cleaned up with Zymo’s ssDNA/RNA Clean & Concentrator kit (Zymo Research D7010) and eluted in 15 μL of molecular biology grade water.

Poly(A)-tailing of cleaned up msDNA was performed by mixing 12.25 μL of molecular biology grade water, 2.5 μL of terminal transferase reaction buffer (NEB B0315), 0.25 μL dATP (of a 10 mM stock diluted from NEB N0440), 3 μL of terminal transferase (TdT, NEB M0315), and 10 μL of msDNA. Reaction was incubated at room temperature for exactly 60 sec and then stopped by heating to 70**°**C for 5 min (∼25 adenosines should have been added in this time).

After this, complementary strand synthesis was performed using a poly(T) primer which also contains an Illumina TruSeq adapter (Supplementary Table 5). The reaction was setup as follows: 28.9 μL of molecular biology grade water, 8 μL of TdT reaction from the previous step, 5 μL of NEBuffer 2 (NEB B7002), 0.1 μL of the poly(T) primer (of a 100 uM stock from IDT), and 5 μL of dNTP mix (of a 10 mM stock from NEB N0447) were mixed and heated to 80**°**C, then allowed to cool to room temperature (facilitates primer annealing). Then, 3 μL (15 units) of DNA Polymerase I, Large (Klenow) Fragment (NEB M0210) was added, and the reaction was incubated at 37**°**C for 30 min. The reaction was purified using the QIAquick PCR Purification Kit (Qiagen 28104) and eluted in 15 uL of molecular biology grade water.

Following this, oligos containing the second Illumina TruSeq adapter (Supplementary Table 5) were annealed by mixing 10 μL of each oligo (top and bottom strand, from a 100 μM stock concentration) with 10 μL of molecular biology grade water and 10 μL of NEBuffer 2, then heated to 95**°**C for 2 min and allowed to cool back to room temperature. 1 μL of the annealed adapters were mixed with 4 μL of the extended msDNA product and 5 μL of Blunt/TA Ligase Master Mix (NEB M0367). This was incubated for 15 min at room temperature, after which the reaction was immediately cleaned up using AMPure XP Beads (Beckman Coulter A63880). Cleanup was performed with 1.8x beads:DNA volume ratio and the product was eluted in 10 uL of molecular biology grade water. Sample was prepared for Illumina sequencing following standard workflows and sequenced on a NextSeq 2000 system. Sequencing reads were trimmed for adapter sequences and aligned to the Retron-Vpa2 operon using Geneious. Visualization of read coverage on the Retron-Vpa2 hyRNA secondary structure was performed using VARNA^70^ (v3-93).

### Split GFP complementation assays

In preliminary experiments developing this assay, a plasmid containing either sfGFP1-10 or SP-sfGFP1-10 with a kanamycin-resistance cassette and a plasmid containing sfGFP11 (either on its own or fused to the SP in the context of Retron-Vpa2 Δnc-hyRNA) with a carbenicillin-resistance cassette were co-transformed into *E. coli* bSLS.114. In all subsequent experiments, a plasmid containing a version of Retron-Vpa2 SP::sfGFP11 with a carbenicillin-resistance cassette and a plasmid containing SP-sfGFP1-10 with a kanamycin-resistance cassette were co-transformed into *E. coli* bSLS.114. For experiments requiring trigger genes, a third plasmid containing the trigger (or an empty vector control) with a chloramphenicol-resistance cassette was also co-transformed. For experiments using Retron-Eco1 to produce msDNA in *trans*, a third plasmid containing the engineered retron with a spectinomycin-resistance cassette was also co-transformed.

For each sample, three individual colonies were picked (biological replicates) and grown until saturation in 0.5 mL LB + antibiotics in a 96-well deep well plate (Thermo Scientific 260251) covered with an air-permeable seal (Breathe-Easier, Diversified Biotech) at 37**°**C with shaking at 1000 rpm. Cultures were then diluted 1:100 in 200 uL of fresh LB + antibiotics in a black, clear-bottom 96-well microtiter plate (Corning 3603). Samples were arranged to have at least one empty well between each set of replicates to avoid bleed-through fluorescence across samples. The plate was covered with an air-permeable seal (Breathe-Easier, Diversified Biotech BERM-2000) and incubated in a microplate reader (Tecan Infinite 200 PRO) with shaking at 37**°**C for 16 hrs with fluorescence measured from the bottom of the wells using an excitation wavelength of 485 nm and emission wavelength of 520 nm. Measurements were taken every 10 min. At the 2 hr timepoint, IPTG and arabinose inducers were added to all cultures. In preliminary experiments developing the assay, inducers were instead added at the 0 hr time point and the experiment was run for 16 hrs. Fluorescence readings were adjusted by subtracting the 0 hr measurement for each sample.

## ACKNOWLEDGEMENTS

Work was supported by funding from the National Science Foundation (MCB 2509382), the Robert J Kleeberg Jr. and Helen C. Kleberg Foundation, and the Gordon and Betty Moore Foundation.

K.Z. was supported by a National Science Foundation Graduate Research Fellowship and a UCSF Discovery Fellowship.

A.C. was supported by a Lundbeck Foundation grant R380-2021-1448.

D.J.Z. was supported by the Fulbright U.S. Student Program, which is sponsored by the U.S. Department of State and the Danish-American Fulbright Commission.

A.D.H was supported by a Ph.D. fellowship from Junta de Castilla y León and European Social Fund Plus (EDU/1868/2022) and an EMBO Scientific Exchange Grant (11561).

G.M. is part of CPR, which is supported financially by the Novo Nordisk Foundation (NNF14CC0001, NNF24SA0098829). This work was also supported by the ERC-AdG 101096548 (INTETOOLS), NNF0024386, NNF17SA0030214, and NNF18OC0055061 grants to G.M, who is a member of the Integrative Structural Biology Cluster (ISBUC) at the University of Copenhagen.

R.P.-R. was supported by a Lundbeck Foundation grant (R347-2020-2346) and a research grant (VIL60763) from VILLUM FONDEN.

S.L.S. is a Chan Zuckerberg Biohub – San Francisco Investigator.

We thank Mylinh Bernardi and Felicia Miller of the Gladstone Genomics Core for their assistance with sequencing co-immunoprecipitated RNA from our pull-down assays.

## AUTHOR CONTRIBUTIONS

Author contributions follow the CRedIT taxonomy (https://www.elsevier.com/researcher/author/policies-and-guidelines/credit-author-statement)

K.Z.: Methodology, Investigation, Validation, Formal Analysis, Writing – Original Draft, Visualization

M.R-M.: Methodology, Investigation, Validation, Writing – Review & Editing

D.P.: Methodology, Investigation, Validation, Writing – Review & Editing

J.M.K.: Methodology, Investigation, Validation, Writing – Review & Editing, Visualization

A.C.: Methodology, Investigation, Validation, Writing – Review & Editing, Visualization

D.J.Z.: Methodology, Software, Formal Analysis, Writing – Review & Editing, Visualization

M.R.M.: Methodology, Software, Formal Analysis, Writing – Review & Editing, Visualization

R.Z.: Methodology, Investigation, Validation

A.D-H.: Investigation

G.M.: Methodology, Writing – Review & Editing, Supervision, Funding Acquisition

R.P-R.: Methodology, Writing – Review & Editing, Supervision, Funding Acquisition

A.G-D.: Conceptualization, Writing – Review & Editing, Supervision

S.L.S.: Conceptualization, Writing – Review & Editing, Visualization, Supervision, Project administration, Funding acquisition

## DATA AND CODE AVAILABILITY

Sequencing data associated with this study are available in the NCBI SRA (PRJNA1320719) https://www.ncbi.nlm.nih.gov/bioproject/PRJNA1320719

Custom code used to process or analyze data from this study is available at: Spacer-seq analysis was performed using code sourced from https://github.com/Shipman-Lab/Spacer-Seq.

## SUPPLEMENTARY MATERIALS

See supplementary files for Supplementary Figures S1-S10 and Supplementary Tables 1-5.

