## Supplementary Figures for "Reverse transcribed ssDNA derepresses translation of a retron antiviral protein"

[1] Gladstone Institute of Data Science and Biotechnology, San Francisco, CA, USA
[2] Department of Bioengineering and Therapeutic Sciences, University of California, San Francisco, CA, USA
[3] Section of Microbiology, Department of Biology, University of Copenhagen, 2100 Copenhagen, Denmark
[4] Structural Molecular Biology Group, Novo Nordisk Foundation Centre for Protein Research, Department of Cellular and Molecular Medicine, Faculty of Health and Medical Sciences University of Copenhagen, Blegdamsvej 3B, Copenhagen, 2200, Denmark
[5] Centro de Investigación del Cáncer, Consejo Superior de Investigaciones Científicas (CSIC), Universidad de Salamanca, 37007 Salamanca, Spain
[6]Chan Zuckerberg Biohub, San Francisco, CA, USA
† These authors contributed equally to this work

**Supplementary Figure 1:** Structural mapping of the escapee mutations in the λ and Stevie exonuclease (*exo*) and recombinase (*beta*/*rec*). **A)** Top view of λ exo in complex with DNA, showing its trimeric assembly (PDB: 3SM4). exo monomers are colored in teal blues, and DNA is shown in red. **B)** Bottom view of the λ exo–DNA complex. Residues I105, C132, and T135 are highlighted as yellow spheres, and two $Mg^{2+}$ ions are shown as green spheres. **C)** Close-up of a λ exo monomer bound to DNA. The position of the DNA 5′ end is labeled. **D)** Structural superposition of Stevie's exo with λ exo bound to DNA. Stevie's protein was aligned to λ exo. Residue E183 is shown as yellow spheres. **E)** Superposition of λ exo–DNA complex (PDB: 3SM4) with the λ exo–beta complex (PDB: 6M9K). Residue E242 in the C-terminal domain of λ beta is highlighted as yellow spheres. **F)** Top view of the helical assembly of λ beta bound to DNA (PDB: 7UJL). **G)** Side view of the λ beta helical assembly with DNA. In panels **(F)** and **(G)**, λ beta is shown as a surface representation, and DNA is colored in red. **H)** Structural superposition of λ beta with the Stevie Rec AlphaFold3 model, revealing a shared N-terminal core. Residues A38, M48, L51, W85, and Q135 are shown as yellow sticks.

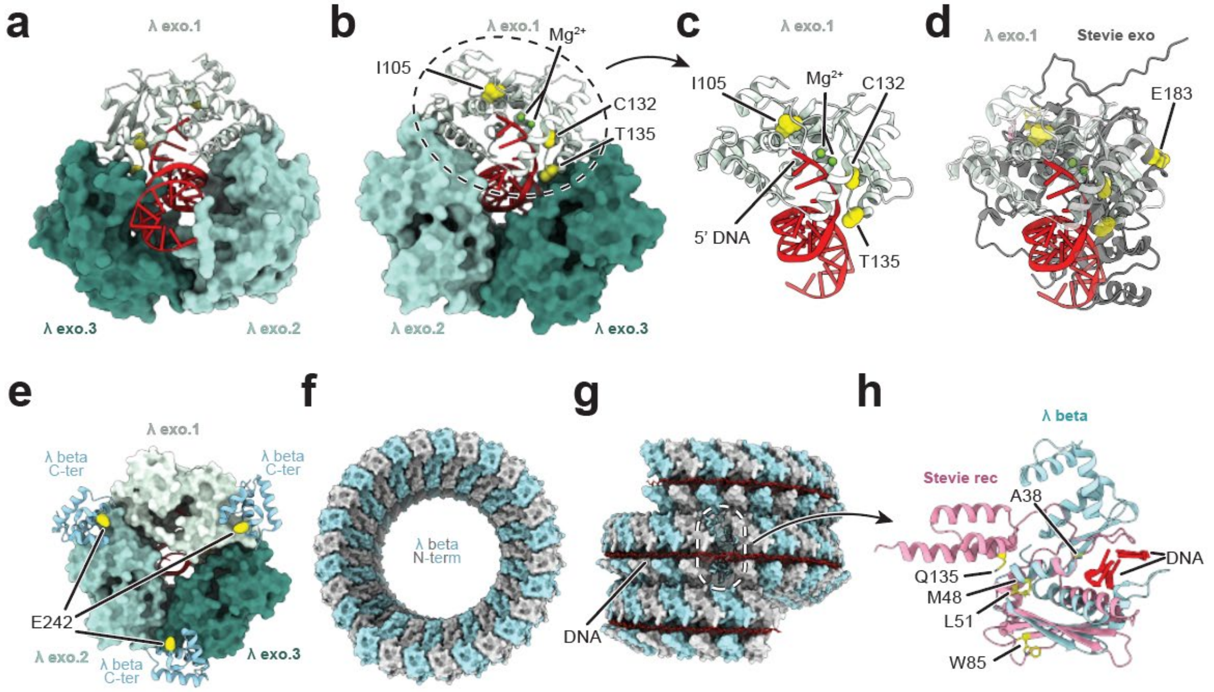

**Supplementary Figure 2:** Growth curves of *E. coli* bSLS.114 strains expressing Retron-Vpa2 with **A)** variants of the lambda Red operon and **B)** individual genes from the lambda Red operon. **C)** Comparison of Retron-Eco6 and Retron-Sen1 growth curves when expressed with the lambda Red operon or Gam. For all plots, solid line indicates the mean and error band indicates the standard deviation of three biological replicates.

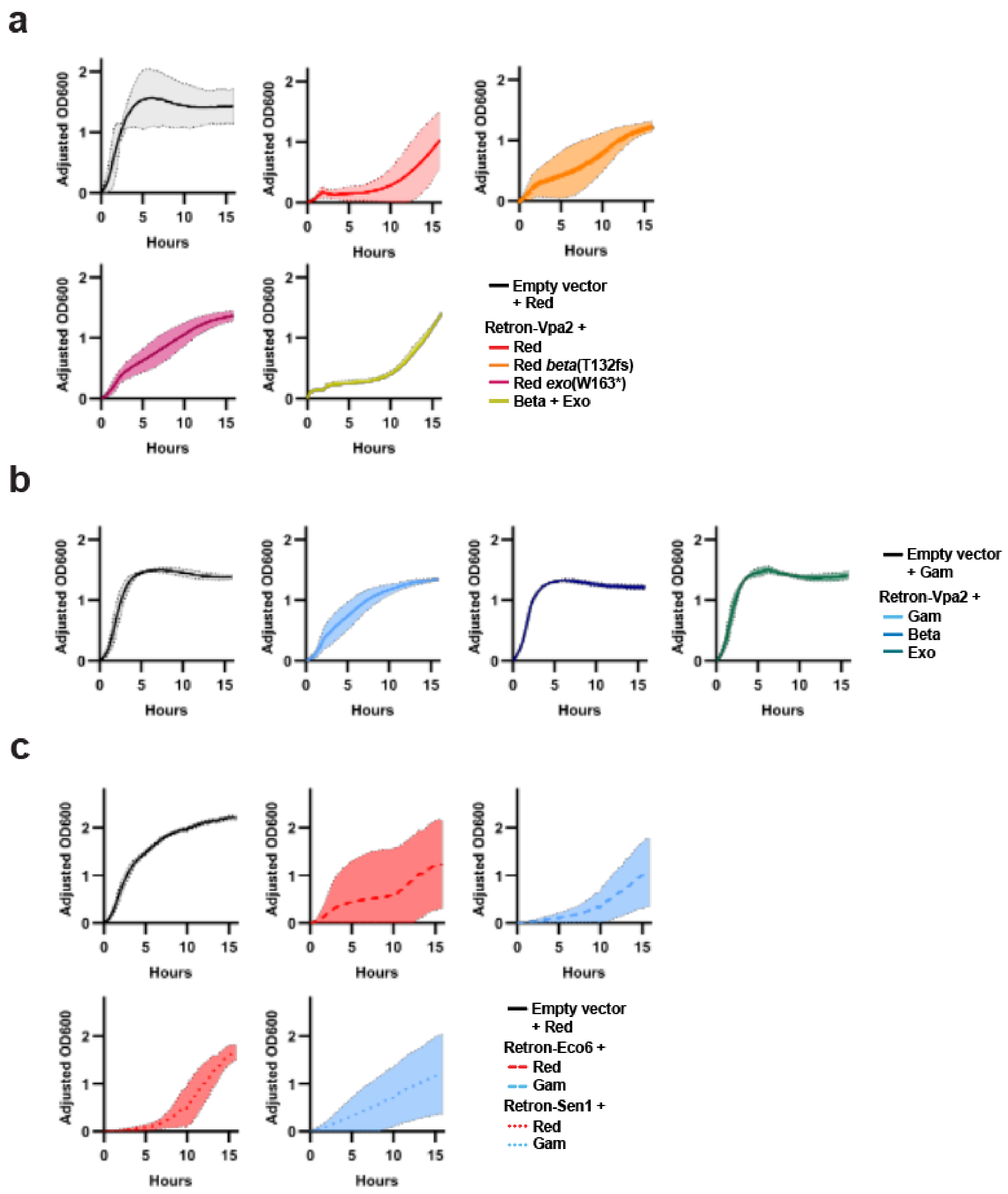

**Supplementary Figure 3:** Coverage plot from RNA-seq of strain expressing Retron-Vpa2 dRT compared to wild-type Retron-Vpa2 (from main text Fig 3a), mapped to the retron operon.

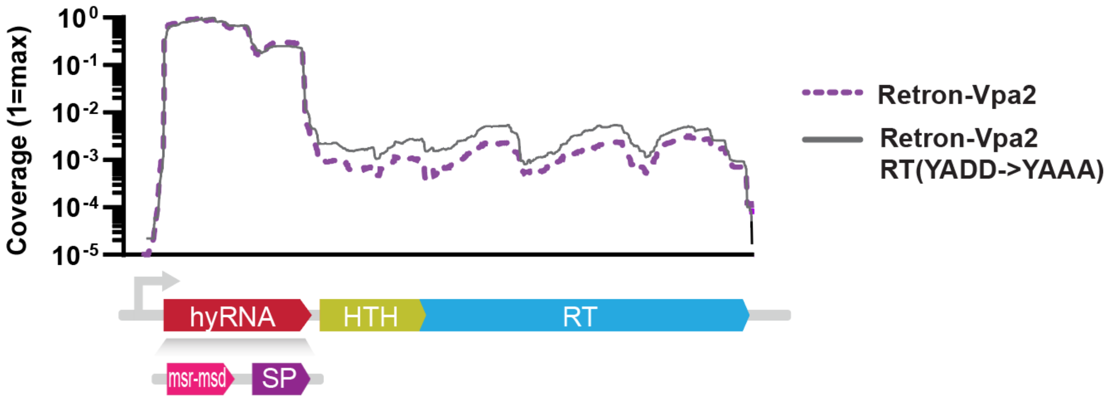


**Supplementary Figure 4: A)** Covariance model for the hyRNA from alignment of 81 Type VI retron homologs. **B)** Consensus sequence of the hyRNA with conserved motifs mapped onto the Retron-Vpa2 hyRNA.

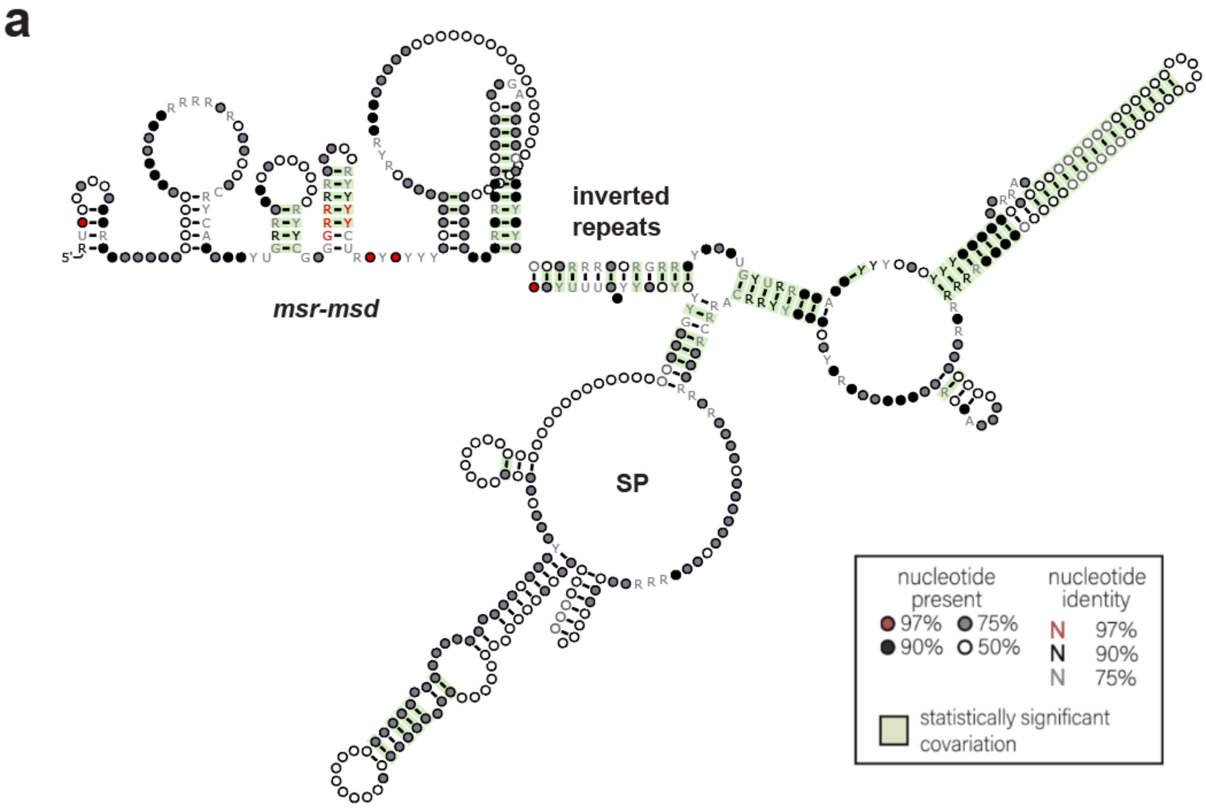

**b**

**Type VI consensus hyRNA:**

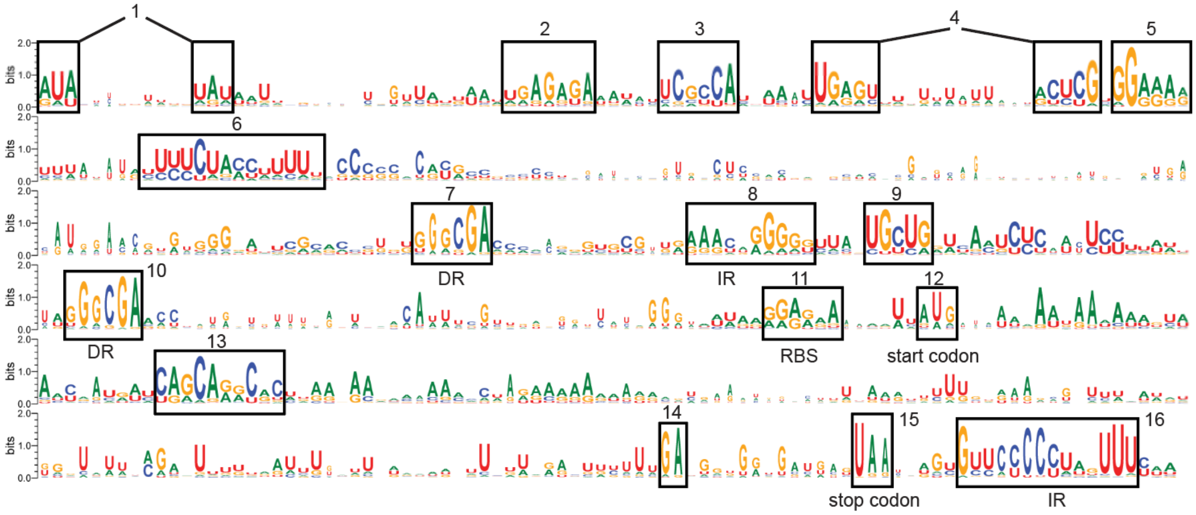


**Retron-Vpa2 hyRNA:**

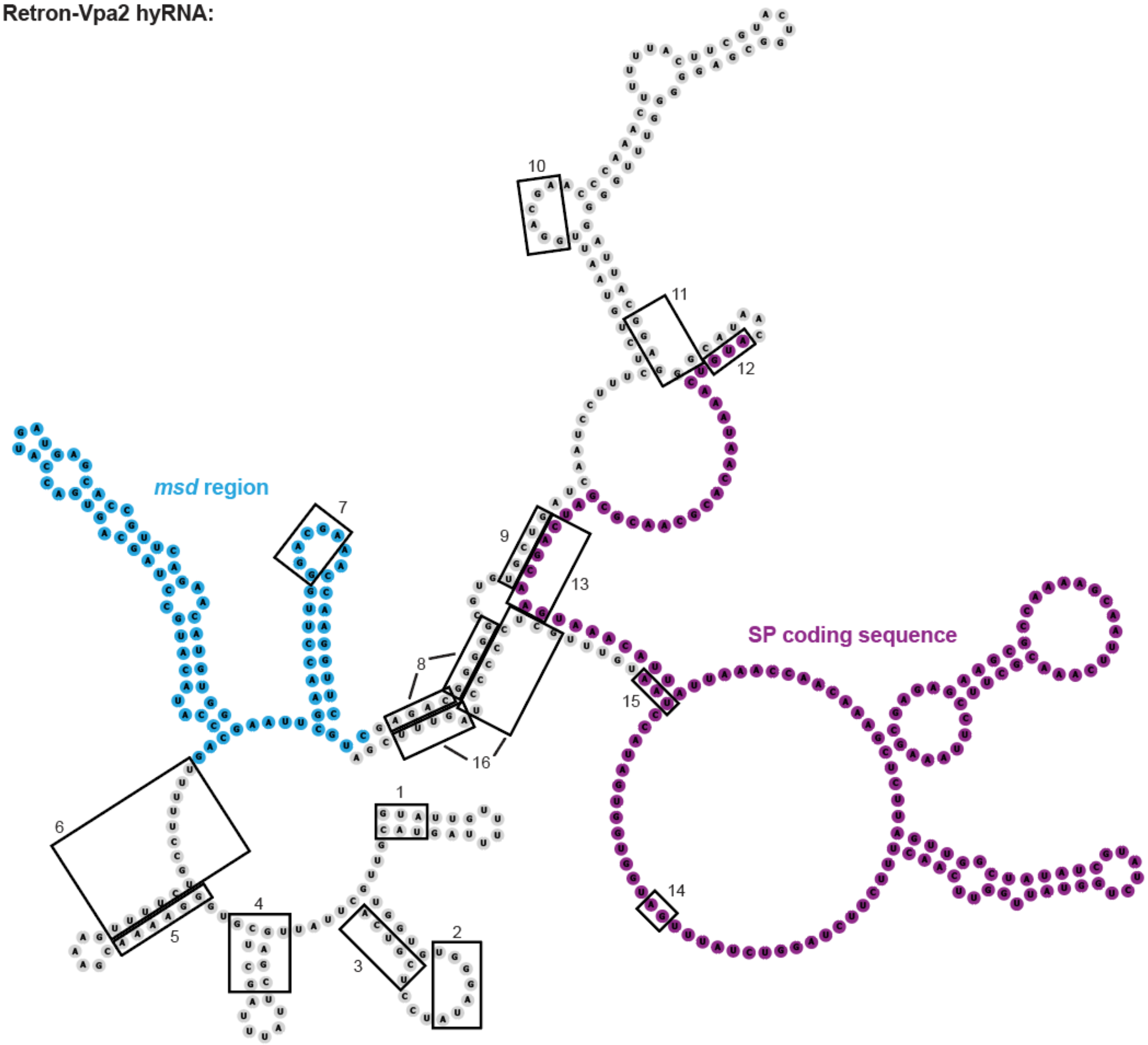

**Supplementary Figure 5:** Structural and evolutionary analysis of Retron-Vpa2 and related small proteins. **A)** AlphaFold prediction of the Vpa2 small protein (SP). Hydrophobic residues I39, F43, and F45 are highlighted in sticks representation. **B)** AlphaFold3 models of the small protein of Vpa2, Eco12, and evolutionary related homologs. Electrostatic potential was plotted on the protein's surface. **C)** Structural superposition of Vpa2, Eco12, and homologous proteins. **D)** Predicted local distance difference test (pLDDT) plotted on the Vpa2 SP alphafold3 model represented as cartoon. **E)** Phylogenetic tree of SP proteins showing evolutionary relationships among Vpa2, Eco12, and related sequences.

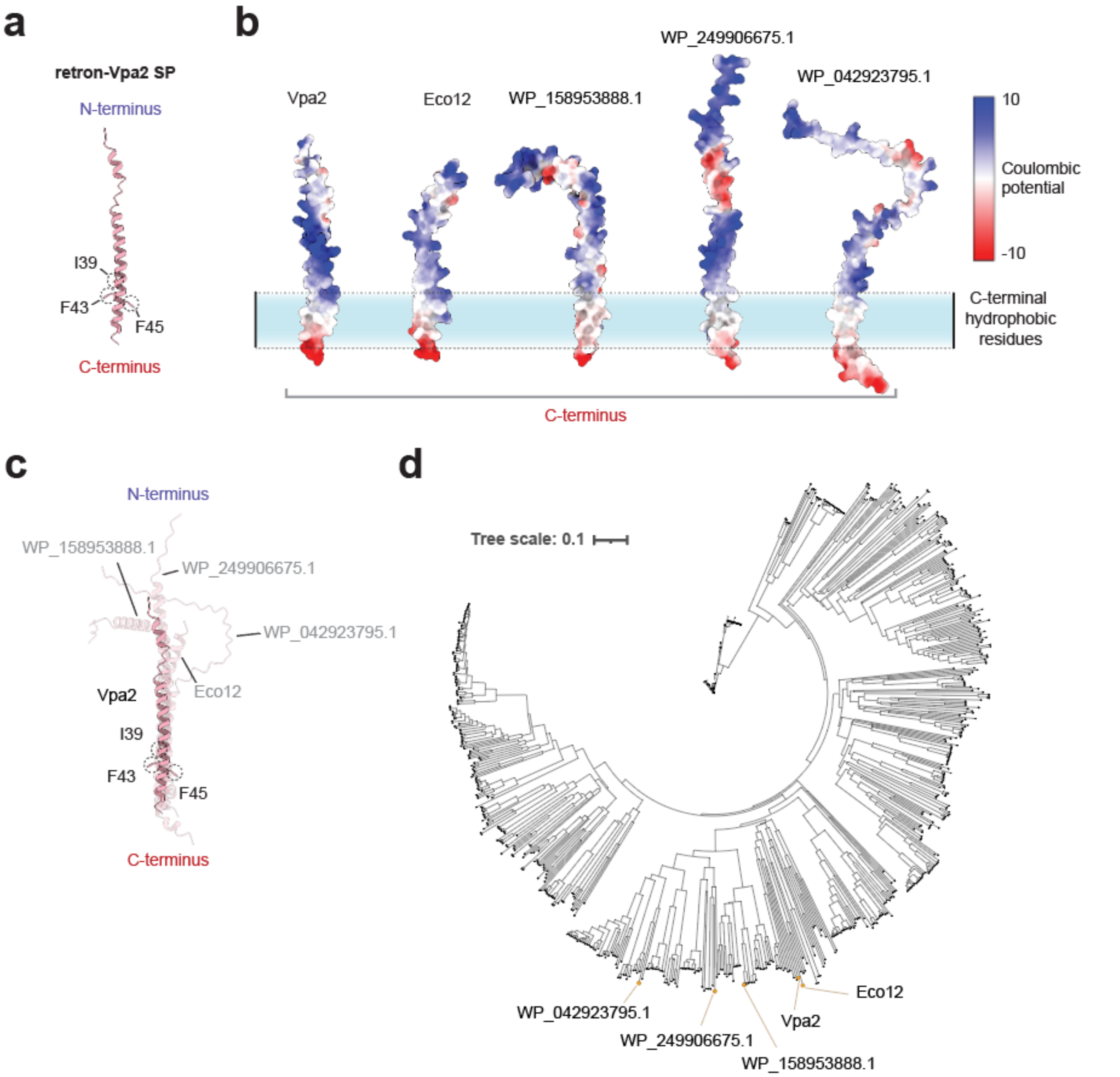

**Supplementary Figure 6:** Growth curves of *E. coli* bSLS.114 strains expressing **A)** variants of the SP effector and **B)** variants of the Retron-Vpa2 operon. For all plots, solid line indicates the mean and error band indicates the standard deviation of three biological replicates.

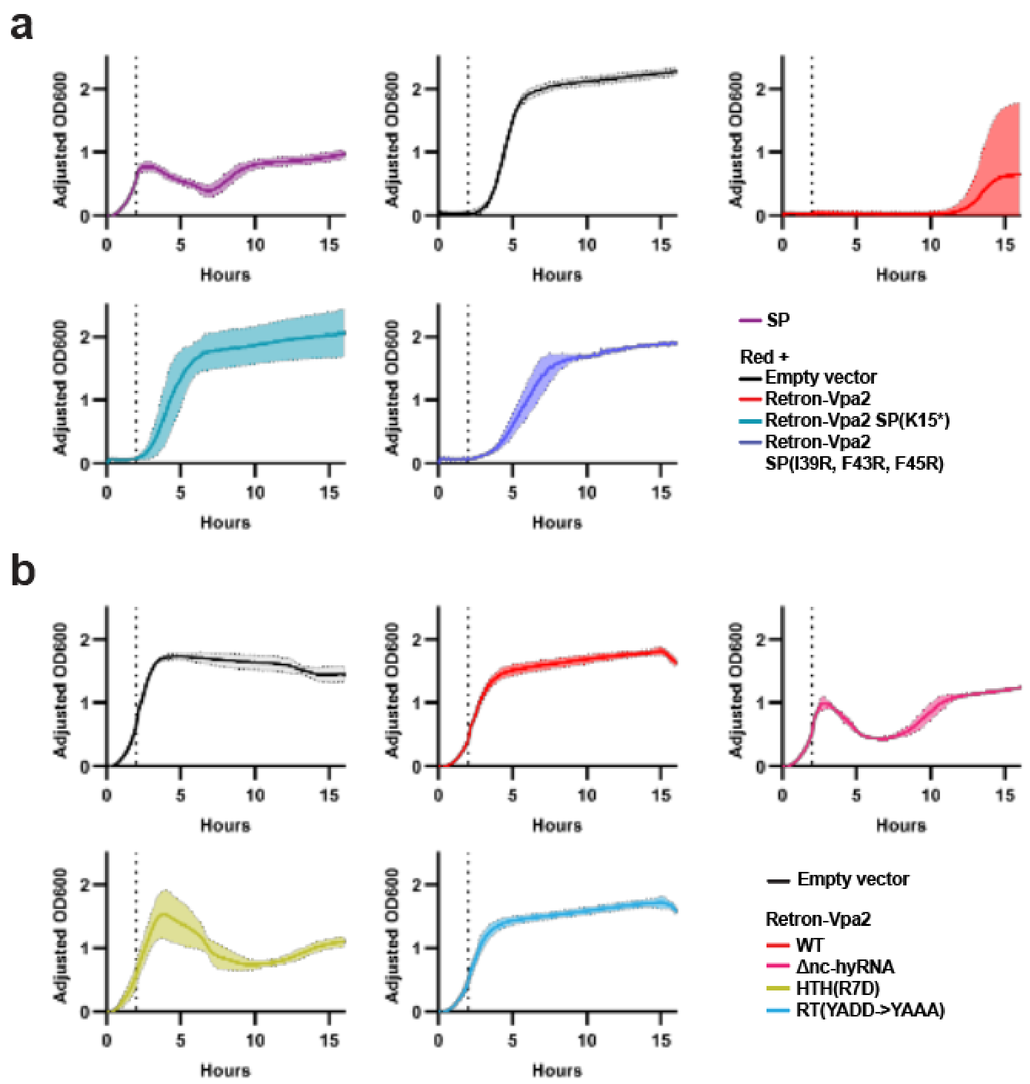

**Supplementary Figure 7:** Uncropped Western blots for HA-tagged HTH and FLAG-tagged RT, as well as PAGE analysis of co-immunoprecipitated RNA, from **A)** pulling down on the RT and **B)** pulling down on the HTH. Dotted boxes indicate regions that were used in the main text Fig 3g-h. **C)** PAGE analysis showing lack of co-immunoprecipitated DNA from both anti-HA and anti-FLAG pull-downs. **D)** Coverage of sequenced co-immunoprecipitated RNA from the RT pull-down mapped to the Retron-Vpa2 operon (normalized to maximum coverage).

**a**

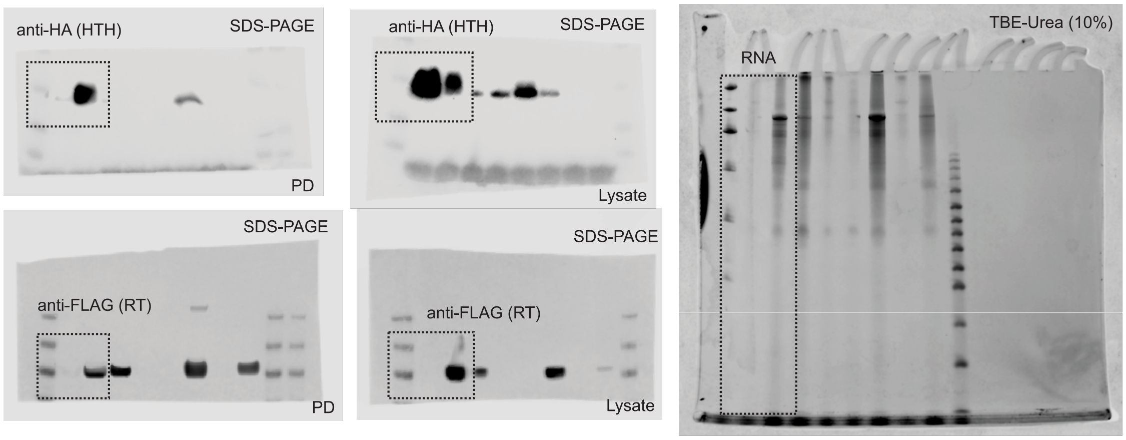


**b**

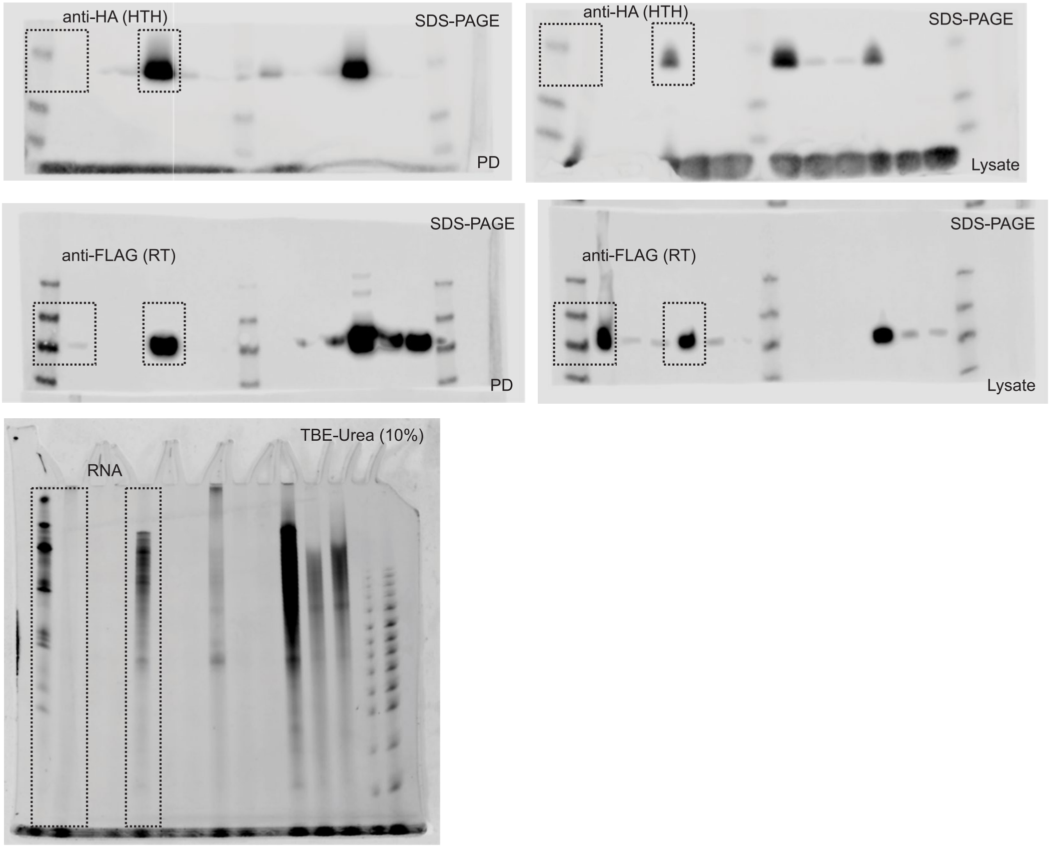

c

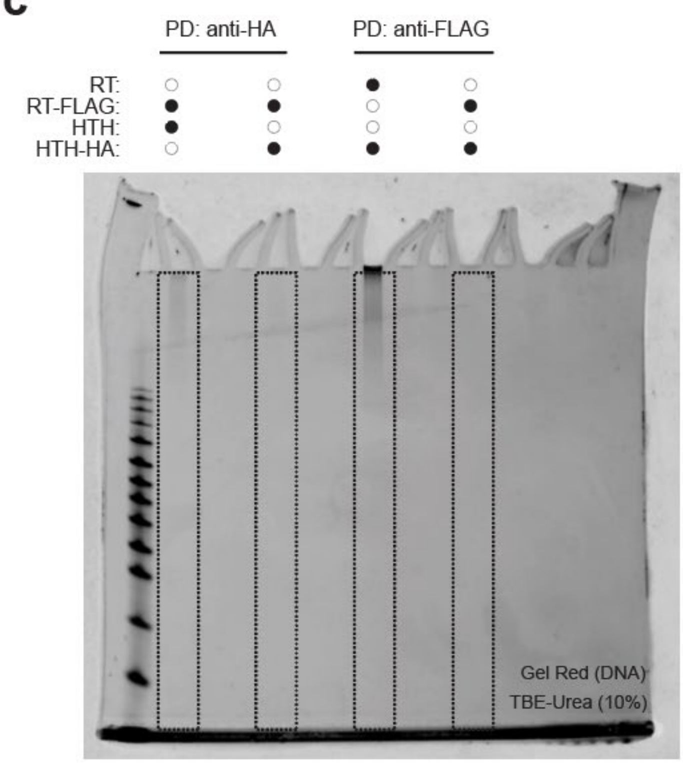

PD: anti-HA
PD: anti-FLAG
RT:
RT-FLAG:
HTH:
HTH-HA:
Gel Red (DNA)
TBE-Urea (10%)


d

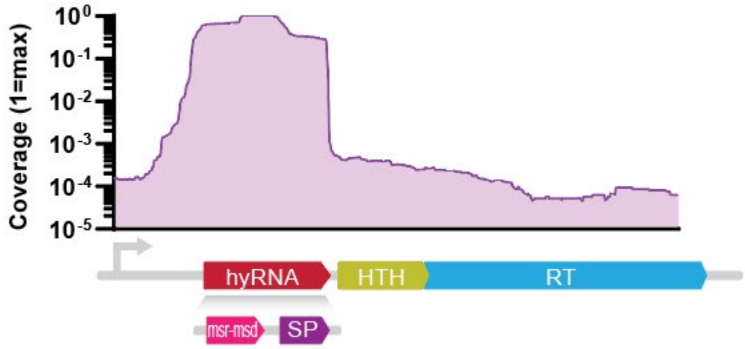

Coverage (1=max)
10^0
10^-1
10^-2
10^-3
10^-4
10^-5
hyRNA
HTH
RT
msr-msd
SP

**Supplementary Figure 8:** Per base coverage (normalized to total spacers acquired) of spacers from *E. coli* bSLS.114 expressing **A)** Retron-Eco6 and **B)** Retron-Sen1, mapped to their respective operons. **C)** Spacer-seq coverage for strains expressing either wild-type Retron-Vpa2 or Retron-Vpa2 dRT. Coverage is mapped across both plasmids in the system and the *E. coli* (BL21) chromosome. The *lacI* gene was excluded from this analysis due to it being present on multiple references.

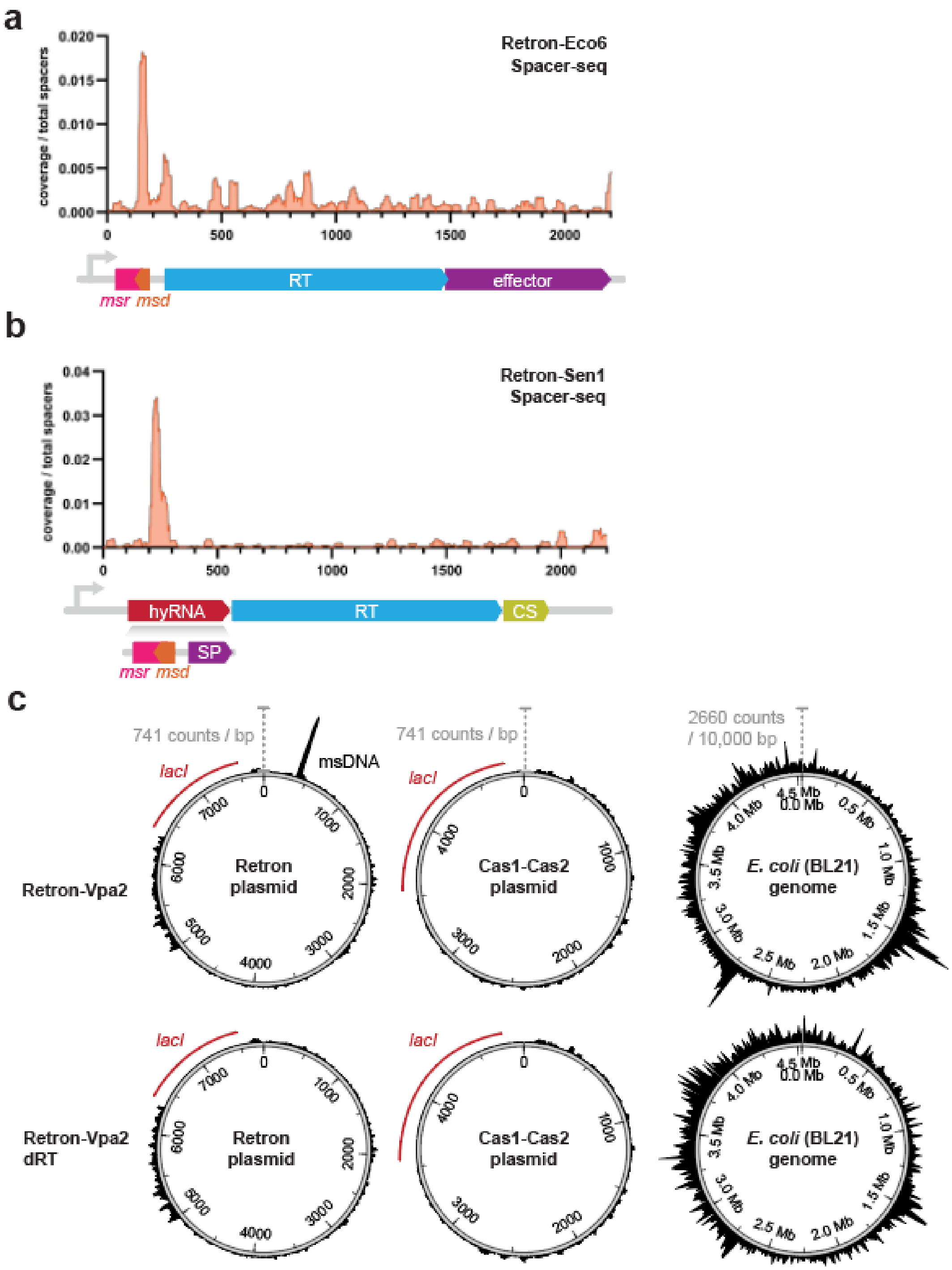

**Supplementary Figure 9: A)** Per base coverage (normalized to total spacers acquired) of spacers aligning to the *msr-msd* region of Retron-Vpa2 for Δ*recB E. coli* bSLS.114 expressing Retron-Vpa2 dSP or Retron-Vpa2 dRT. Shown for comparison is Retron-Vpa2 dSP in *E. coli* bSLS.114 (same as in main text Fig 4d-h). **B)** Uncropped gel of Retron-Vpa2 msDNA bands. Dotted boxes indicate regions that were used in the main text Fig 4i.

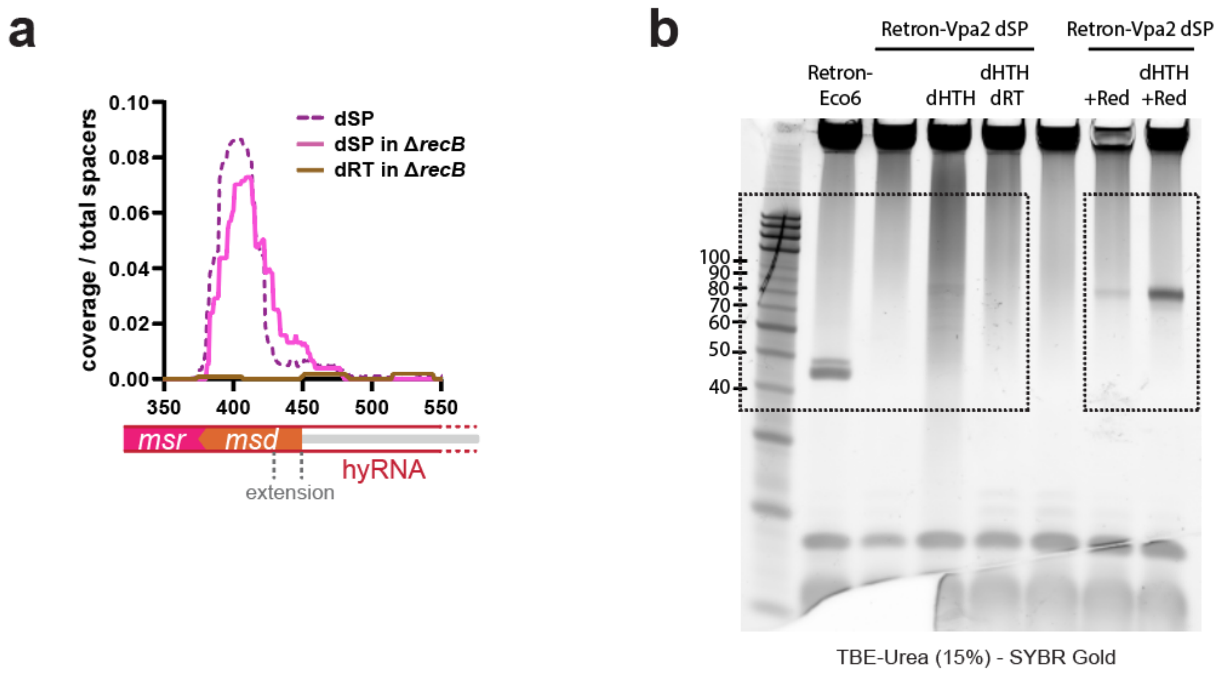


**Supplementary Figure 10:** Growth curves of *E. coli* bSLS.114 strains expressing variants of a split sfGFP system (see main text Fig 5b). Solid lines indicate the mean and error bands indicate the standard deviation of three biological replicates.

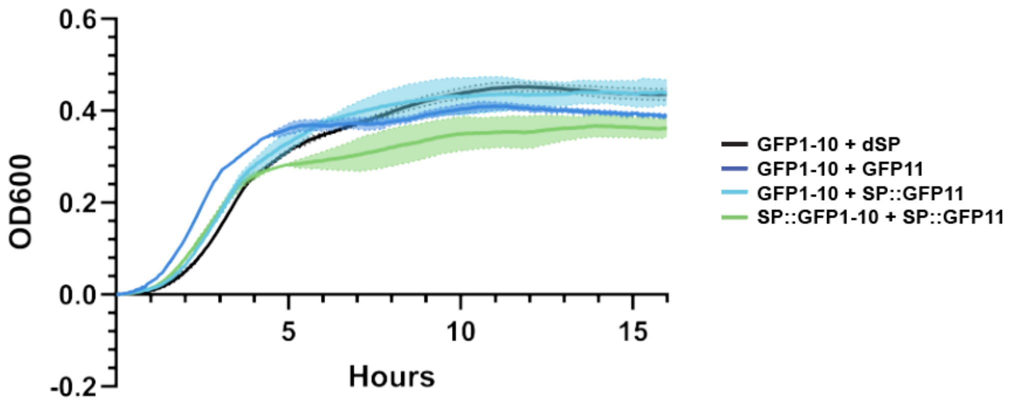
